# Exploring Biosurfactant-Producing Bacteria from Waste-Contaminated Sites near Dhaka City

**DOI:** 10.64898/2026.03.18.712685

**Authors:** Ummey Fatema Tun Amina, Maliha Mahzabin, Sabrina M Elias

## Abstract

Industrial waste containing hydrophobic pollutants, like oils and hydrocarbons, is toxic and difficult to degrade, posing both ecological and human health risks. Biosurfactants are eco-friendly surface-active compounds produced by microorganisms, known for their ability to lower surface and interfacial tension, enhancing the solubility and bioavailability of hydrophobic compounds, facilitating their breakdown. The current study focuses on isolating biosurfactant-producing bacteria from industrial waste sources near Dhaka, Bangladesh, and characterizing their properties, determining potential usage. Using diesel-enriched nutrient agar, bacterial strains were isolated and screened for biosurfactant production by oil displacement, emulsification index (E24%), and drop collapse assay. The most promising isolates were characterized according to their biochemical activities and 16S rRNA amplicon-based sequencing. Isolation and characterization of the surfactants have been carried out using chromatographic techniques. The identified bacteria passed the drop collapse and oil displacement tests. CTAB agar assay, indicates their anionic nature, showing an emulsification index ranging 10-41%. The potential biosurfactant producers belong to *Bacillus, Pseudomonas, Acinetobacter, and Enterobacterium* genera. The surfactants showed antibacterial, antifungal, and plant growth promotion activity and have been characterized in terms of pH stability, salinity, adhesion, and temperature tolerance. The study successfully identified and characterized potential biosurfactant-producing bacteria from industrial waste, highlighting their efficiency in breaking down hydrophobic pollutants and hydrocarbons. These microorganisms provide a green and economical substitute for synthetic surfactants due to their biodegradability and lower toxicity. Upon further research and scaling, these bacteria can be a good source of biosurfactants for potential applications in industrial, agricultural, and biomedical fields.

**IMPORTANCE:** The study carries high significance as it creates multi-disciplinary scopes for utilizing these environmentally adapted biosurfactant-producing bacteria in industry, agriculture, and medicine. Since the bacterial isolates have hydrocarbon degradation ability, upon optimization for higher production, industrial usage in oil refinery and other industries can be adopted. Due to their biodegradable nature, usage in wound healing bandages and as antimicrobial agents in medicine will be noteworthy. The isolates have plant growth promotion ability and utilizing them as biofertilizer will reduce the dependency on chemical fertilizers. This is the first detailed report on biosurfactant-producing bacteria from this industrial waste-polluted Turag River of Dhaka City. Moreover, it compiles detailed screening protocols and methods for analyzing such environmentally friendly microbes. Such characterization also opens the scope for optimizing the production of the surfactant compounds on a large scale and utilizing them commercially.

## INTRODUCTION

Industrial waste is typically composed of hydrocarbon garbage, heavy metals, and other pollutants, which are mostly hydrophobic in nature. Due to their water-repelling nature, these pollutants are not easily degradable and pose a health risk to humans and other elements in the environment. To remove such pollutions, synthetic surfactants are used regularly, which are amphiphilic in nature, i.e, they can interact with both hydrophobic and hydrophilic compounds. Thus, surfactants aid in waste removal by emulsifying hydrophobic pollutants, enhancing their solubility and facilitating their breakdown or removal from contaminated environments. However, these synthetic surfactants are often toxic, non-renewable, and environmentally persistent, raising concerns about their long-term sustainability (Banat et al. 2010) As an alternative to them, biosurfactants are eco-friendly due to their biodegradability and low toxicity. Biosurfactants are microbial origin surface active compounds that contain both hydrophilic and hydrophobic moieties. These compounds are produced by various organisms, including fungi, bacteria, and actinomycetes. These amphiphilic molecules are primarily located on microbial surfaces but can also be secreted extracellularly. They effectively reduce surface and interfacial tension, enhance emulsification, and break down the hydrophobic pollutants as well. Thus, they can be used in bioremediation, industrial processes, oil recovery, pharmaceuticals, and agriculture. Also, they often show anti-bacterial, antifungal, and anti-adhesive properties, which are beneficial for different medical applications (Gudiña et al. 2010).

Mostly biosurfactants are produced by the genera Bacillus, Pseudomonas, Acinetobacter, Rhodococcus, etc. Based on their chemical structure and their microbial origin, these surfactants are typically assigned to different classes. Major classes of biosurfactants include phospholipids, glycolipids, lipopeptides, polymeric biosurfactants, etc. Bacillus spp. produces lipopeptides such as surfactin, fengycin, and iturin, which have surface-active and antimicrobial properties (Saiyam et al. 2024). Among them, surfactin is one of the most potent biosurfactants currently recognized (Arima et al. 1968). The glycolipids rhamnolipids, sophorolipids, and trehalolipids are the most well-known types (Md 2012). Pseudomonades are notable for producing rhamnolipid, widely used in bioremediation due to their emulsification efficiency and ability to degrade hydrocarbons. The chemical composition of biosurfactants is classified into various categories, including fatty acids, polymeric and particulate biosurfactants, glycolipids, glycolipopeptides, glycoproteins, and lipopeptides. Surfactin, a common lipopeptide, is synthesized through non-ribosomal biosynthesis, which is catalysed by surfactin synthetase, a large multienzyme peptide synthetase complex. Other lipopeptides like iturin, lichenysin,, arthrofactin, etc are synthesized by similar enzyme complexes. Non-ribosomal peptide synthetases (NRPSs) play a crucial role in lipopeptide biosynthesis, while the rhl quorum-sensing system is essential for glycolipid biosurfactant production in *Pseudomonas* species. Biosurfactant composition and emulsifying activity are dependent on both the producer strain and the culture conditions; consequently, the type of carbon and nitrogen source, as well as the C:N ratio, nutritional limitations, chemical and physical parameters like pH, aeration, temperature, and divalent cations affect both the quantity of the manufactured biosurfactant as well as the kind of polymer produced (Salihu et al. 2009).

Biosurfactants produced by microbes increase its ability to grow, spread, and persist in challenging environments. It helps them access hydrophobic nutrients, move across surfaces, and face the stress. These compounds also give them advantages by disrupting competitor microbes and improving root colonization. But not only for the microbe itself, but biosurfactants are useful for bioremediation of locations contaminated with hydrocarbons, oil recovery, heavy metals, and pesticides (Datta and Chattopadhyay 2024). Surfactants are being reported for use in a wide range of industrial domains, including environmental protection, agriculture, petroleum, food, and the pharmaceutical industry (Deleu and Paquot 2004; Rostás and Blassmann 2009)((Singh et al. 2007) (Moldes et al. 2011). They serve as a biocontrol agent to shield plants against a variety of diseases, increasing agricultural production (Moldes et al. 2011; Simpson et al. 2011). Biosurfactants are superior to chemical surfactants in terms of biocompatibility (Mohan et al. 2006), low toxicity (Desai and Banat 1997), biodegradability, accessibility of the basic materials, and stability at extreme pH and temperature (Muthusamy et al. 2008). According to Mulligan (2005), a high-quality surfactant may reduce the interfacial tension between hexadecane and water from 40 to 1 mN/M and the surface tension of water from 75 to 35 mN/m ((Muthusamy et al. 2008) (Makkar and Cameotra 1997). Temperatures as high as 50°C, a pH range of 4-5 to 9.0, a 50g/l concentration of NaCl, and a 25g/l concentration of Calcium did not impact the lichenysin that the *Bacillus licheniformis* strain produced (Muthusamy et al. 2008). Their functional properties indicate that they have far greater potential for application in agriculture, particularly in plant protection and soil management techniques that aim to improve the environment to produce nutritious food. Hence, studies on the characteristics of surfactants and the search for bacteria that produce biosurfactants are intriguing fields of study. A deeper understanding of genetic regulatory mechanisms could facilitate the development of metabolically engineered strains with the capability of enhanced production characteristics.

Microorganisms growing in pollutants and waste often gain adaptability to survive the pollutants, developing unique metabolic capabilities. Biosurfactant production by microbes is one such example that helps them survive in hydrocarbon-rich and heavy metal-polluted habitats. Urban waste disposal sites, industrial effluents, and oil-contaminated soils are rich in such microbial diversity and can be an excellent source of potential biosurfactant-producing bacteria. Tongi, near the Dhaka metropolitan area, is one of the most industrialized and polluted regions in Bangladesh that generates large amounts of industrial waste. The presence of naturally adapted biosurfactant-producing bacteria in such environments poses opportunities for biotechnological exploitation. The underexplored microbial diversity and their adaptation mechanism need attention for systematic exploration, potential scalability for industrial applications, and ensuring their eco-friendly multidimensional usage in our daily life.

In this study, we have explored biosurfactant-producing bacteria from the industrial waste-contaminated water sample near Dhaka Metropolitan City. The workflow of the study is depicted in Figure 1. The isolates were screened for their biosurfactant-producing ability using the oil displacement test, emulsification index (E24%), and drop collapse assay. The strains were characterized using different biochemical tests, unveiling their metabolic and enzymatic activities as well as 16s rRNA amplicon sequencing-based genus-level identification. Thin-layer chromatography was used to characterize the nature of the surfactant, followed by the extraction of the surfactant compound. The stability of the biosurfactant-producing ability under varying pH, temperature, and salinity conditions was assessed to evaluate their application and adaptability in industry. Additionally, the antibacterial, antifungal, adhesion, biofilm formation, and plant growth promotion ability of the bacterial strains were explored to evaluate their application in the medical and agricultural fields. By identifying efficient biosurfactant-producing bacteria from a pollutant environment, the study shows a cost-effective, environmentally friendly alternative to synthetic surfactants. However, to maximize their potential, an increase in the yield of biosurfactant molecules needs to be achieved.

**Figure 1:**
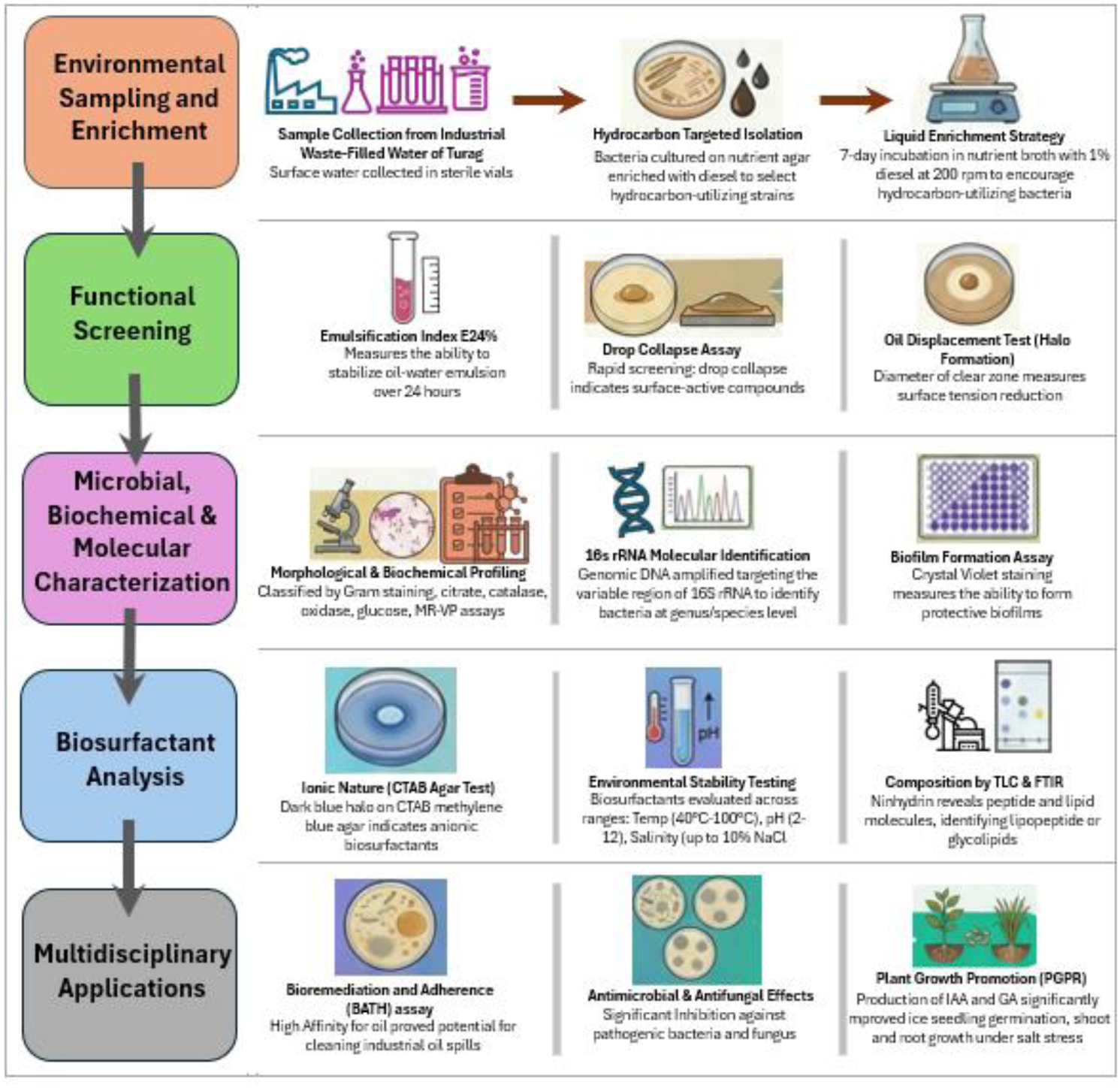
Schematic Diagram of the workflow

### 2. MATERIALS AND METHODOLOGY

### 2.1 Sample Collection

Samples of surface water were taken from the Tongi Khal, near the industrial zone of Tongi, a city close to Dhaka, Bangladesh. The samples were taken in sterile vials, sent right away to the Independent University Bangladesh’s laboratory, and kept there at 4°C pending additional examination.

### 2.2 Growth media preparation

Nutrient broth and agar medium were employed to separate the bacteria that produced the biosurfactants. The liquid medium composition was peptone (10.0 grams), meat extract (10.0 grams), sodium chloride (5.0 grams), per liter of distilled water without adding any agar. The solid media was made by adding 1.5% of the agar, adjusting the pH to 7.2 ± 0.2, and autoclaving the mixture for 15 minutes at 121°C and 15 pounds of pressure.

### 2.3 Inoculum preparation

The spread plate method was used to plate the samples on the nutrient agar medium after they had been serially diluted. The samples were incubated for 24 hours at 37°C inside an aerobic environment. Each petri dish contained diesel as the hydrocarbon source, and a control without diesel was additionally maintained with the assistance of cotton buds. Hydrocarbon sources were added to the medium to enrich it. In a 250 ml Erlenmeyer flask with 1% liquid-soluble fraction of diesel, these 24-hour-grown colonies were infected with 100 ml of nutrient broth medium at 200 rpm and 30°C for seven days (Dunlap and Beckmann 1988). Once the incubation phase had lasted for one week, 1 milliliter of the inoculum was moved to 99 milliliters of nutritional broth that contained 1% diesel and left for two days.

### 2.4 Screening of the isolates for biosurfactant production

For thirty minutes, the culture medium was centrifuged at 3000 r.p.m (revolutions per minute). The cells were disposed of, and the supernatant was collected. Due to the extracellular production of the biosurfactant, the cell-free extract was obtained by centrifugation and utilized for subsequent examination (Femi-Ola et al. 2015).

#### 2.4.1 Oil Displacement Test

When a potential surfactant-containing solution was added to an oil-water interphase, the diameter of the clear zone could be measured using an approach called oil displacement (Morikawa et al. 2000). The biosurfactant’s effectiveness in reducing surface tension can be determined by measuring its diameter. This test involved filling a 90 mm-diameter petri dish with 15 milliliters of distilled water. The water surface received 100 µl of diesel, while the oil surface received 20 μl of cell culture supernatant. After 30 seconds, the circumference and the distinct halo seen in visible light were measured (Rodrigues et al. 2006). A drop of water served as the negative control.

#### 2.4.2 Emulsification Index (E24%) measurement

The Cooper and Goldenberg method was used to determine the biosurfactant’s emulsification capacity towards diesel and kerosene (Cooper and Goldenberg 1987). One milliliter of cell-free extract and two milliliters of hydrocarbon were combined in a test tube following the centrifugation of the sample culture, and the mixture was homogenized by vortexing for two minutes. After 24 hours, the emulsion activity was examined, and the emulsification index (E24%) was computed by multiplying the total height of the liquid layer by the total height of the emulsion layer, followed by multiplication by 100. As a positive control, SDS with E24% = 72 was used to compare the outcomes.

#### 2.4.3 Drop Collapse Assay

The experiment was conducted following instructions provided by Jain et al. (1991). After a minute, a drop of the culture supernatant was carefully placed on a glass slide that was already coated with oil and examined. The presence of biosurfactant, i. e, the positive result, is indicated if the supernatant drop collapses and disperses across the oil-coated surface. If the drop persisted for a minute, it was considered as negative. As a control, an experiment with distilled water was also carried out. This experiment was carried out in a 96-well microtiter plate lid, where each well of the lid was filled with 2 μL mineral oil and was kept for 1 hour at room temperature. After this, 5 μL of the bacterial culture supernatant was added to the surface of the oil. The drop shape was visually inspected after 1 minute. Production of flattened, collapsed drops was considered positive for biosurfactant production, indicating surface activity. On the other hand, stable and round-shaped drops indicated the absence of biosurfactant activity.

### 2.5 Biochemical and Molecular Characterization

#### 2.5.1 Gram Staining

Gram staining was carried out to differentiate the Gram-positive and Gram-negative bacteria based on the thickness of their peptidoglycan layer. A bacterial smear was heat-fixed on a slide, stained with crystal violet, followed by iodine treatment, decolorization with ethanol, and counterstaining with safranin. The results were observed under a microscope.

#### 2.5.2 Biochemical Tests for metabolic and enzymatic characterization

A series of biochemical tests was carried out following standard microbiological procedures to determine the metabolic and enzymatic activities of the isolates.

##### 2.5.2.1 Motility, Indole and Urease (MIU) test

To assess if the bacteria are motile and whether they can produce indole and urease, the MIU test was carried out, where the isolates were inoculated in the MIU medium by a single stab with a sterile needle and incubated at 37 °C for 24 hours. Motility was confirmed by observing bacteria growing away from the stab line, forming turbidity. Urease activity was confirmed by observing a pink-red color, as the presence of urease will break down urea, releasing ammonia, ultimately turning the phenol red indicator in the medium pink red. Indole production was determined by the formation of a red ring at the surface after the addition of Kovac’s reagent, indicating breakdown of tryptophan to indole

##### 2.5.2.2 Kligler Iron Agar (KIA) test

In KIA agar, the bacterial culture was inoculated by stabbing followed by streaking and was incubated at 37 °C for 24 hours. Representative color changes and the presence of gas or H_2_S were observed. In the test tube, a yellow slant and butt indicate fermentation of both glucose and lactose, a red slant and yellow butt indicate fermentation of glucose only. H_2_S production is indicated by black precipitation, and gas production can be detected by observing any presence of cracks or bubbles in the agar.

##### 2.5.2.3 Citrate Utilization Test

Bacterial Isolates were streaked onto Simmons’ citrate agar slants and incubated at 37°C for 24-48 hours. Positive citrate utilization was indicated by a colour change from green to blue, which confirms their ability to use citrate as a sole carbon source.

##### 2.5.2.4 Starch Hydrolysis Test

Bacterial cultures were streaked onto starch agar plates and incubated at 37°C for 24 hours, followed by flooding with iodine solution. A clear zone around the bacterial growth indicated positive starch hydrolysis, confirming production of amylase as opposed to the absence of a clear zone with blue-black coloration, indicating negative starch hydrolysis.

##### 2.5.2.5 Oxidase test

A fresh bacterial colony was smeared on a filter paper strip soaked with N,N,N′, N′-Tetramethyl-1,4-phenylenediamine. A positive oxidase test was indicated by a color change to dark purple within 30 seconds, suggesting the presence of cytochrome c oxidase.

##### 2.5.2.6 Catalase test

A loopful of respective bacterial culture was taken on a glass slide and 3% Hydrogen peroxide was added as a drop. The presence of a gas bubble indicates a positive catalase reaction, since catalase can break down hydrogen peroxide and produce oxygen and water.

##### 2.5.2.7 Methyl Red (MR) and Voges-Proskauer (VP) test

Bacterial culture was inoculated in MR-VP broth followed by incubation for 48 hours at 37°C. For the MR tests, methyl red indicators were added in 5 drops. A red colour indicated a positive MR test, which means stable acid has been produced from glucose fermentation. For the VP test, Barritt’s A (α-naphthol) and Barritt’s B (KOH) solutions were added to the incubated bacterial culture in MR-VP broth. A reddish colour development within 15-30 minutes indicated a positive VP test, suggesting acetoin production.

#### 2.5.3 16s rRNA-based sequencing

To identify the powerful strain at the molecular level, 16S rRNA analysis was carried out. With the use of a ProFlex PCR system, 16S rRNA gene was amplified from the genomic DNA using the universal forward primer 27F (5′AGTTTGATCCTGGCTCAG3′) and reverse primer 1492 R (5′AAGGAGGTGATCCAGCC3′). The PCR tube consists of ∼50 ng of DNA template, 1X PCR master mix, 10µM of forward and reverse primers adjusted with nuclease-free water. In the PCR protocol, denaturation was carried out for five minutes at 95°C, annealing for one minute at 57°C, and extension for seven minutes at 72°C. This was followed by thirty-five cycles of temperature cycling. Electrophoresis on 1% (w/v) agarose gel was used to isolate the amplified 16S rRNA gene. Sanger sequencing was carried out in the Genetic Analyzer 3500 by Applied Biosystem from Thermosphere Scientific. Serial dilution was performed to retrieve the pure culture. The sequences were aligned with the NCBI 16S rRNA database, and the sequence showing the highest similarity index was considered as the identity of the unknown bacteria. However, for further confirmation of the culture, 16S rRNA amplicon-based next-generation sequencing was performed in a Miseq Illumina platform targeting the 16SrRNA V3 and V4 region (Klindworth et al. 2012) with 300bp reads. Qiime2 and DADA2 were used to process the reads, and species-level taxonomic classification was performed using the silva138 database.

#### 2.5.4 Biofilm formation assay

The experiment was started by adding a small number of isolates to 3–5 milliliters of nutrient broth, which was then placed in an incubator set at 37°C for 24 hours. The first well, known as the negative control, received 100μL of distilled water. Fifty microliters of each isolates were then added to the other wells. After sealing the plate, it was left to incubate at 37°C for the entire day. Each well’s cells were cleaned three times using distilled water. Following a 10-minute staining process with 125μL of 0.1% crystal violet, each well was rinsed with distilled water and allowed to dry. In each well, 200μL of ethanol was added, and after 15 minutes of mixing, the optical density (OD) at 600 nm was measured (Sharma and Kaur 2015).

### 2.6 Characterization of the surfactant compound

#### 2.6.1 Ionic characterization via CTAB agar test

According to Siegmund and Wagner (1991), the creation of a dark halo zone in a plate containing Cetyl Trimethyl Ammonium Bromide (CTAB) and methylene blue was evaluated to determine the generation of extracellular biosurfactant. 0.005 g of methylene blue and 0.2 g of CTAB were added to BH agar (Bushnell-Huss Agar) medium supplemented with 1.8% agar, and the mixture was sterilized at 121 °C for 15 minutes. After adding 10 µl of the culture supernatant to the plate, it was incubated at 30°C for 48 hours. Formation of a dark blue halo surrounding the culture would indicate an anionic nature of the biosurfactant due to ion matching with the cationic CTAB-methylene blue agar complex (Saravanan and Vijayakumar 2012).

#### 2.6.2 Emulsion stability studies

Emulsion stability tests were conducted in accordance with Campos et al.’s instructions to ascertain the optimal circumstances for the stability and functionality of the biosurfactant generated by isolates (Campos et al. 2019). The E24% experiment (using diesel) was used to determine the emulsion stability at various temperatures, pH levels, and salt concentrations. By completing the E24% assay of cell-free culture with diesel and incubating the emulsion created at 40°C, 80°C, and 100°C, the impact of temperature on the stability of the emulsion was ascertained. By bringing the cell-free broth’s pH (using 1N HCl or 1N NaOH) or salt concentration (by adding NaCl) to the appropriate level and computing E24% indices after 24 hours, the impact of pH and salinity on the stability of the emulsion was ascertained. Salt concentration ranged from 0% to 10% (2%, 6%, and 10%), whereas pH ranged from 2 to 12 (2, 7, and 12).

#### 2.6.3 Thin layer Chromatography

The biosurfactant-producing isolates were cultured in 100 ml of BHM (Bushnell-Huss medium) broth, which was supplemented with 2% (w/v) glucose, 2% (v/v) molasses, and 2% (v/v) oil. This culture was kept with agitation (2 × g) at 30°C for 7 days. Cell-free supernatants were generated by centrifuging at 5000 × g at 4◦C by removal of bacterial cells. This cell-free supernatant was then treated with 2N HCl for 16h at 4°C so that the solution had an acidic pH of 2.0. The solution was then centrifuged at 5000 × g for 10 min at 4°C, and the resulting precipitate was collected. This acid precipitate fraction (APF) was then dissolved in 100mg/mL water, and the pH was adjusted to 7.0 using 1N NaOH (Rani et al. 2020). The CFS sample and clean BHM culture medium were stored at −20°C until further use. Thin-layer chromatography (TLC). was used to separate the CFS and APF fractions. Before use, the TLC plates were heated to 120°C for 30 minutes to activate the silica plates. A 70:26:4 ratio of chloroform: methanol: water was used as a solvent system. 100mg of each CFS and APF was separated on activated TLC plates using 400 µL of chloroform: methanol (2:1, v/v).

Chloroform: methanol: water (70:26:4) was used as the developing solvent system for the separation of 100 mg of each of CFS and APF in 400 µL of chloroform: methanol (2:1, v/v) on activated TLC plates. Plates were examined in UV light (λ = 254 nm), and the retention factor (Rf) values were computed by dividing the sample’s travel distance by the solvent’s travel distance. The dry plates were sprayed with a 0.25% (w/v) ninhydrin in acetone solution and incubated at 115 °C for five minutes to detect peptides (Xia et al. 2014).

### 2.7 Additional properties of the bacterial strains

#### 2.7.1 Bacterial Adhesion to Hydrocarbons (BATH) assay

Cell viability in the presence of oil was examined using the BATH assay. Centrifugation was used to collect the overnight-grown culture in broth, and it was then resuspended three times in phosphate buffer (g l−1, 16.9 K_2_HPO4, and 7.3 KH_2_PO4). At 610 nm, the optical density was determined to be approximately 0.5. After vortexing 2 milliliters of cell suspension with 100 microliters of oil for three minutes, the aqueous layer was separated. The oil coating was removed, and the optical density was measured following one hour of undisturbed incubation (van der Vegt et al. 1991). The following formula (Equation 1) was used to determine the % adherence to oil:

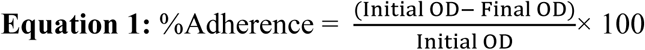

Where initial OD refers to the optical density of the cell suspension in buffer solution before mixing with oil. And final OD *(also known as OD_Shaken_)* refers to the optical density of the aqueous phase after the cell suspension and the oil have been mixed and allowed to separate. Thus, it represents the bacterial cells remaining in the aqueous phase that did not adhere to the oil.

#### 2.7.2 Agar Diffusion assay to test antimicrobial activity

Using the Burkholder agar diffusion assay, the strains were further examined for their antibacterial efficacy against a broad panel of clinical and environmental bacterial species (Burkholder et al. 1944). To reach mid-log phase and a cell density of 106 CFU mL−1, all target bacterial strains and experimental isolates were cultured in LB broth at 27°C for 16–18 hours with constant shaking. This was determined using standard curves that connected plate counts on LBA plates to optical density at 600 nm (OD_600_). After centrifugation at 4°C for 20 minutes at 6500 × g, the cells were suspended in sterile deionized water. Each bacterial suspension (80 µL) was then combined with 7 mL of melted half-strength LB agar, and the mixture was transferred into 15mm diameter culture plates. The Plates were incubated at 24°C after an aliquot (15 µL) of experimental isolates was placed in the middle of the bacterial lawn. After 24 to 48 hours, zones of bacterial growth inhibition next to the spotted isolates’ inoculum were seen. The diameter of these zones varied between 3 and 6 mm, based on the efficacy of the isolates’ antimicrobials that permeated into the agar. To verify the results, each trial was run three times, and the assay was done twice overall (Xia et al. 2014).

#### 2.7.3 Antifungal activity

Antifungal activity of biosurfactant extracts was evaluated using the agar-well diffusion method. Potato Dextrose Agar (PDA) was used as the medium in a plate, and four wells were generated 3 cm away from the center of the plate. At the center, a 0.3cm diameter mycelial plug of the fungus was inoculated. In the control well 50 µL of methanol was placed, and in the rest of the wells 50µL of the biosurfactant solutions were loaded. Here, we used two types of biosurfactant solutions. One is the cell-free supernatant, and the other is the filtered version of the cell-free supernatant.

We also assessed the antifungal activity of the biosurfactant inoculum using the spread plate method. 3 mL of the biosurfactant inoculum was evenly spread onto the surface of each Potato Dextrose Agar (PDA) plate. A fungal mycelial plug (approximately 3 mm in diameter) was aseptically placed at the center of each plate. The plates were incubated at room temperature (25 ± 2°C) for a specified period to assess fungal growth inhibition. Activity was assessed against plant-pathogenic fungi such as Colletotrichum sp. and Pestalotiopsis, as well as various non-pathogenic fungal species.

#### 2.7.4 Plant Growth Promotion Activity

##### 2.7.4.1 Rice seedling stage screening upon application of bacteria

Bacterial culture was prepared using overnight growth of the bacteria in Nutrient broth (NB) at 37 °C. Using 1% (v/v) sodium hypochlorite, rice seeds were surface-sterilized and washed thrice with sterile water to remove traces of chemicals. Then the seeds were treated with O/N grown bacterial culture to observe the changes in germination and growth. We used two experimental groups here, one where the seeds were soaked in 2 mL of bacteria for 2 hours, shaking, and in the other, the seeds were immersed in 2 mL bacterial culture along with 150mM 2 mL NaCl solution. The seeds were then moved to sterile petri dishes with 4mL sterile water, and the moisture condition was maintained for 7 days. A similar experiment was carried out, submerging the seeds in 2 mL of cell-free supernatants to observe the effects of the surfactant. Seed germination percentage was tracked every day while incubating them at 28°C. The percentage of germination was calculated as the number of germinated seeds divided by the total number of seeds multiplied by 100 (Equation 2).

Seedling growth measurements were carried out after 7 days. Root length (cm) and Shoot Length (cm) were measured using a calibrated scale. Vigor index was determined by multiplying the germination percentage by seedling length (Equation 3).

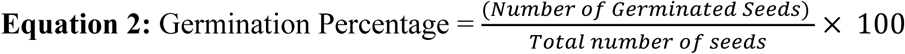

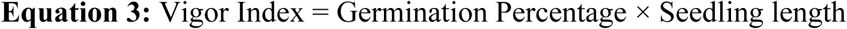

##### 2.7.4.2 Indole Acetic Acid (IAA) production assay

The 10-day-old bacterial culture was centrifuged to obtain cell-free supernatants, and indole-3-acetic acid (IAA) production was assessed. For this colorimetric assay, 2mL freshly prepared Salkowski reagent was combined with 1mL of the supernatant. Salkowski reagent contained 0.5 M FeCl_₃_ in 35% perchloric acid (Gordon and Weber 1951). The mixture was then incubated at room temperature for 30 minutes in the dark, allowing the development of color. Presence of IAA was confirmed with the appearance of a pink to reddish color (Patten and Glick 2002). Color intensity was recorded through photography for documentation. For quantitative assessment, a standard curve was generated ranging from 0 to 50μg mL⁻¹ using commercially available IAA diluted in sterile distilled water. Absorbances were taken at 530nm using a UV–Visible spectrophotometer (Shimadzu UV-1800). Estimation of the sample concentration was done based on the calibration curve (Patten and Glick 2002).

##### 2.7.4.3 Detection of Gibberellic Acid (GA)

Gibberellic acid (GA) detection was carried out using the 2,4-dinitrophenylhydrazine (DNPH) colorimetric method. Briefly, both GA standard and sample solutions were taken in 1 mL quantities to a tightly stoppered Pyrex test tube. A control containing solvent without GA was also prepared. In each tube, 1 mL DNPH reagent was added and incubated at 30 °C for 1 hour, allowing formation of hydrazone. Then 5mL of 10% potassium hydroxide (KOH) in 80% methanol was added to each tube, the solution was mixed and allowed to stand for 5 minutes for color development. The appearance of a wine-red color indicated the presence of GA. The reaction mixture was then diluted with distilled water, and absorbance was measured at 430 nm and 540 nm using a spectrophotometer with the blank as reference. The measured absorbance values were assumed to be proportional to the GA standard concentration. DNPH assay is based on the formation of colored complexes between gibberellins and DNPH under alkaline conditions (Davies 2004).

### 2.8 Extraction of Biosurfactant

One of the best-performing isolates (S2), 1% diesel, and nutritional broth were used to prepare the bacterial inoculum. It is incubated for 120 hours at 37°C. The supernatant is then separated from the medium by centrifugation for 20 minutes at 6,000 rpm at 37°C. Following separation of the supernatant, 1N HCl was used to adjust the pH to 2.0. The supernatant was treated with 2:1 ethanol: chloroform mixture to extract the biosurfactant. At temperatures 40°C - 45°C and 100-200 millibars, a rotary evaporator was used to separate the biosurfactant (extract) from the solvent at the bottom. For additional biosurfactant characterization, the volume of biosurfactant generated by the bacterial consortium is measured and kept at -10°C (Samadi et al. 2007).

## 3. RESULTS

### 3.1 Hydrocarbon-Utilizing Bacterial Isolates found in industrially polluted water

From the water sample collected from Tongi Turag Khal, contaminated with industrial waste, six bacterial cultures were isolated in nutrient agar culture medium enriched with diesel as a hydrocarbon source. The names IUBFASM-tgbS1, IUBFASM-tgbS2, IUBFASM-tgbS3, IUBFASM-tgbS4, IUBFASM-tgbS5, and IUBFASM-tgbS6 were assigned to the six isolates. Later, the bacteria were tested for their ability to grow with kerosene-enriched media as well, which showed better results in some cases. Diesel and kerosene are complex mixtures of hydrocarbons that consist mainly of alkanes (paraffins), cycloalkanes (naphthenes), and aromatic hydrocarbons.

### 3.2 Isolated bacterial strains showed biosurfactant-producing ability

Biosurfactant-producing ability of the isolates was confirmed using the standard tests for assessing oil displacement, emulsification, and drop collapse activities.

#### 3.2.1 Oil Displacement Test Showed Surface Tension Reduction

When a surfactant-containing solution is added to an oil-water interphase, the diameter of the clear zone can be measured using a technique called oil displacement. The diameter evaluation makes it possible to determine how well a particular biosurfactant reduces surface tension (Jain et al. 1991). Amphiphilic compounds, such as biosurfactants, possess both hydrophilic and hydrophobic areas, and they are surface active, which enables them to attract and repel water. Because of their special structure, they can interact with oil and water, lowering the tension that exists between the two phases’ interfacial regions. Hence, biosurfactants are useful in reducing the interfacial tension between water and oil in the setting of oil displacement, which enhances wettability and maximizes oil recovery. For the oil displacement test, the isolates gave positive results, shown in Figure 2. In the oil displacement test IUBFASM_tgbS6 showed the highest value, and IUBFASM_tgbS2 showed the lowest value, which is 6 cm^2^ and 2.7 cm^2,^ respectively. Thus, it implies that the isolated bacteria were able to replace oil on the water’s surface by creating biosurfactants.

**Figure 2:**
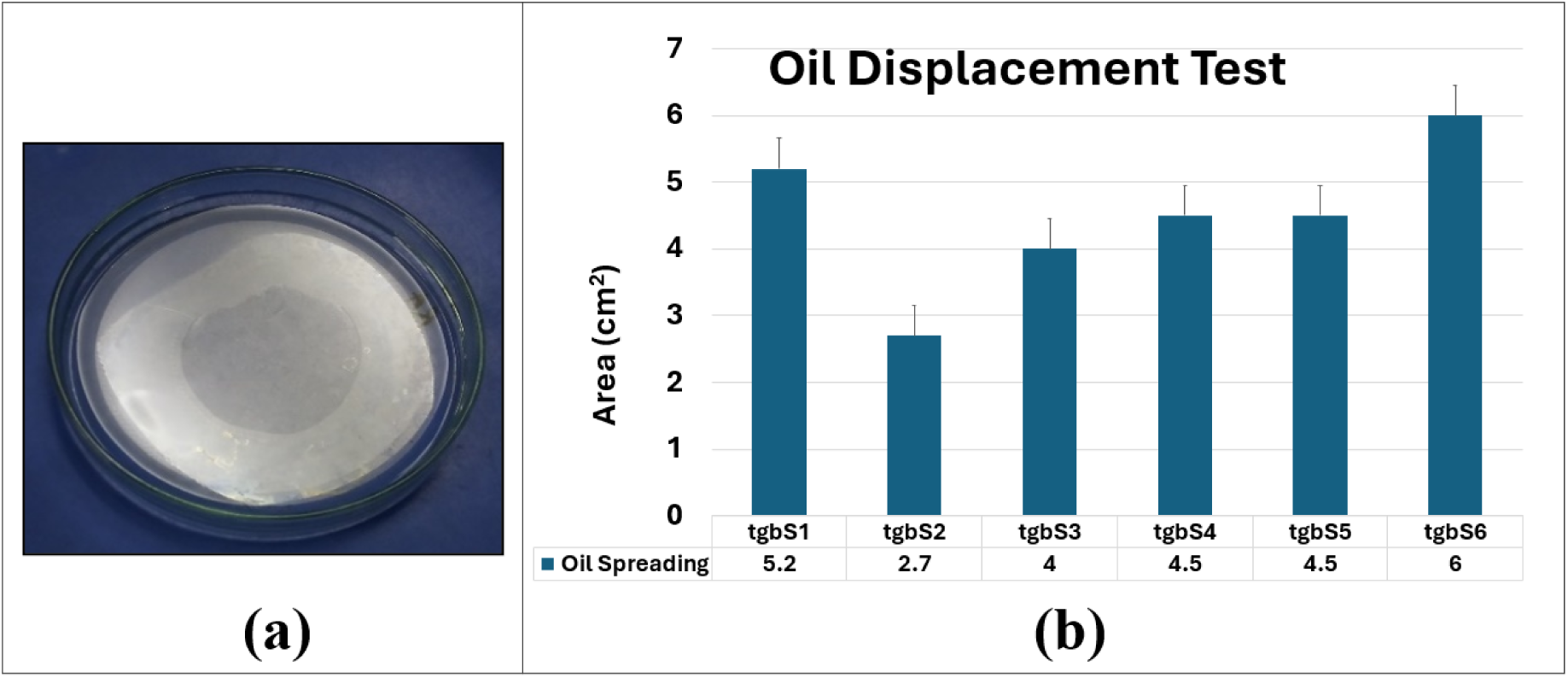
(a) Oil displacement test showing the presence of a clear zone or halo in IUBFASM_tgbS6 (b) Oil Displacement ability of the six isolates in terms of area (cm^2^)

#### 3.2.2 E24% Highlights Strain-Specific Emulsification Ability

The emulsification index (E24%) was utilized to quantify the emulsification activities of the biosurfactant generated by the bacterial consortia that break down polycyclic aromatic hydrocarbons. Improving the interaction of oil and water is one of the roles of biosurfactants, where they form and stabilize emulsions between immiscible liquids. By measuring the emulsification index, biosurfactants can be found to be more effective at improving the contact between water and oil. An elevated emulsification index indicates a high level of oil-water interaction and stronger emulsifying property. The emulsification index (E24%) of the six isolates in diesel and kerosene is shown in Figure 3 and Supplementary Figure 1. IUBFASM_tgbS1 and IUBFASM_tgbS6 showed high emulsification indices in both diesel and kerosene; however, variation was observed in the emulsification capacity of bacteria for diesel and kerosene, e.g. IUBFASM_tgbS2 showed higher E24% in diesel but lower in kerosene.

**Figure 3:**
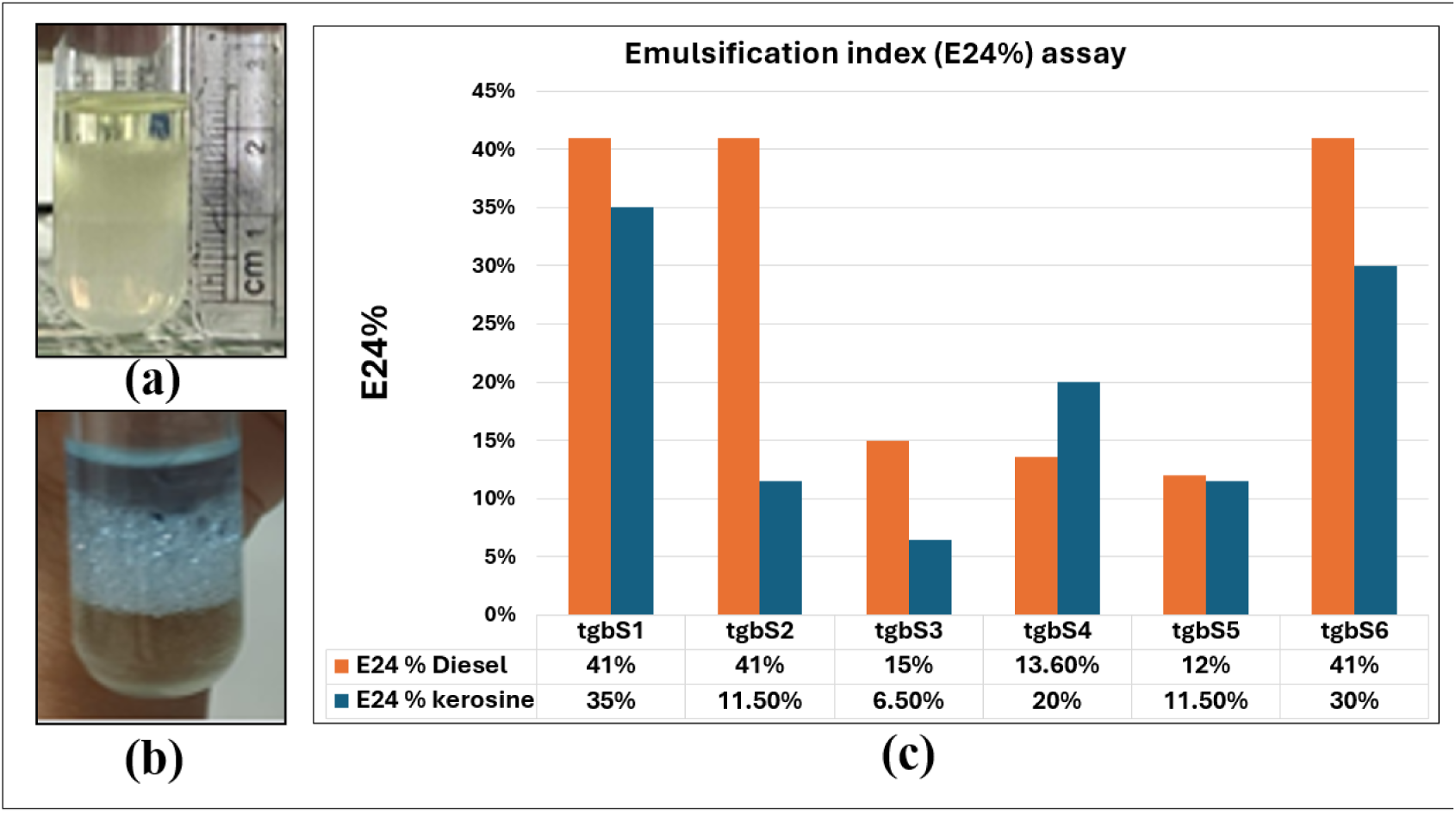
Emulsification Assay (a) Emulsification Assay of IUBFASM_tgbS6 in Diesel (a) and Kerosene (b). Emulsification Index of all isolates (c)

#### 3.2.3 Drop collapse assay reveals reduction of surface tension

In the drop collapse assay, if the drop of supernatant collapsed after one minute and spread throughout the oil-coated surface, it is indicated that biosurfactant is present (positive). If the drop continued beyond a minute, it was noted as negative (Jain et al. 1991). When separated culture supernatants are applied to a solid surface coated with oil, the droplets expand or even collapse due to a decrease in the force or interfacial tension between the liquid drop and the hydrophobic surface (Ali and Rajak 2013). All six isolates showed positive results from the drop collapse test, indicating the effectiveness of the generated surfactant to quench the oil.

### 3.3. Biochemical and Molecular characterization of the bacterial isolates

#### 3.3.1 Gram Staining Uncovers the Diversity of Bacterial Strains

The isolates (IUBFASM_tgbS1, IUBFASM_tgbS3, IUBFASM_tgbS4, and IUBFASM_tgbS6) were found to be rod-shaped, gram-positive (+) according to Gram Staining, indicating the presence of a strong peptidoglycan layer on their cell walls, which is characteristic of Gram-positive species. On the other hand, IUBFASM_tgbS2 and IUBFASM_tgbS5 were found as Gram-negative bacteria.

#### 3.3.2 Metabolic and Enzymatic Activities Confirmed by Biochemical Tests

##### Indole production

The biochemical result for the 6 isolates is shown in Table 1. It was found that IUBFASM_tgbS5 can convert tryptophan into indole, a property shared by *E. coli* and other bacteria (Whitman and Parte 2009). Negative (-) for the others, which indicates these isolates are not indole producers (Versalovic 2011).

**Table 1:**
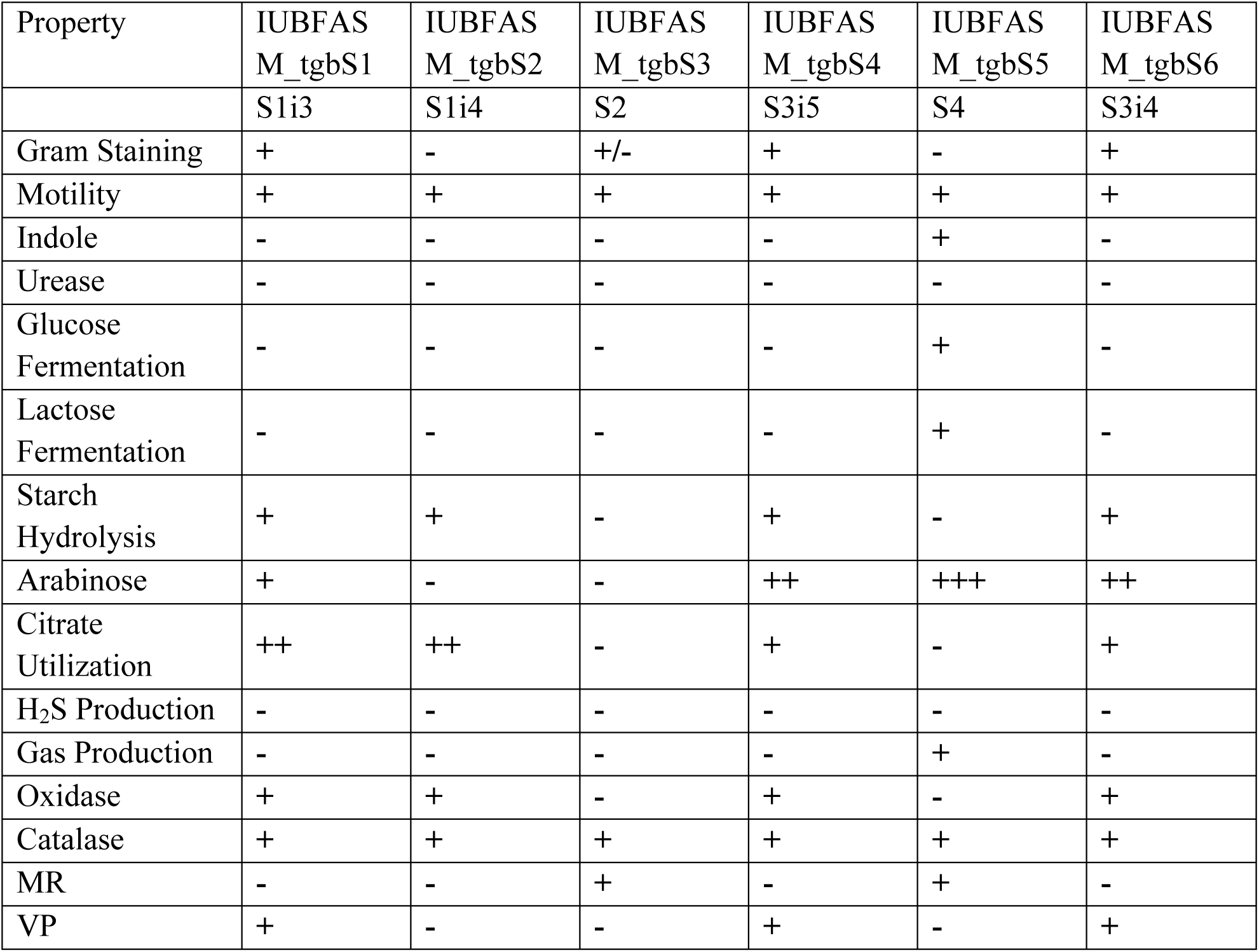
Biochemical Test Results of the six biosurfactant-producing isolates.

| Property | IUBFAS<br>M_tgbS1 | IUBFAS<br>M_tgbS2 | IUBFAS<br>M_tgbS3 | IUBFAS<br>M_tgbS4 | IUBFAS<br>M_tgbS5 | IUBFAS<br>M_tgbS6 |
| --- | --- | --- | --- | --- | --- | --- |
|  | S1i3 | S1i4 | S2 | S3i5 | S4 | S3i4 |
| Gram Staining | + | - | +/- | + | - | + |
| Motility | + | + | + | + | + | + |
| Indole | - | - | - | - | + | - |
| Urease | - | - | - | - | - | - |
| Glucose Fermentation | - | - | - | - | + | - |
| Lactose Fermentation | - | - | - | - | + | - |
| Starch Hydrolysis | + | + | - | + | - | + |
| Arabinose | + | - | - | ++ | +++ | ++ |
| Citrate Utilization | ++ | ++ | - | + | - | + |
| H<sub>2</sub>S Production | - | - | - | - | - | - |
| Gas Production | - | - | - | - | + | - |
| Oxidase | + | + | - | + | - | + |
| Catalase | + | + | + | + | + | + |
| MR | - | - | + | - | + | - |
| VP | + | - | - | + | - | + |

##### Motility and Urease test

All isolates showed motility, meaning they have structures for mobility, such as flagella. All of them were found to be negative (-) in the urease test result. Hence, urea cannot be hydrolyzed by any of the isolates to provide CO₂ and ammonia (Garcia 2010).

##### Carbohydrate fermentation

The glucose fermentation process was positive (+) for IUBFASM_tgbS5 only, and other biosurfactant-producing bacteria were found to be negative after 24-hour incubation. Fermenting glucose allows it to produce acid and potentially gas. Also, the presence of β-galactosidase is shown by IUBFASM_tgS5’s ability to ferment lactose (Versalovic 2011). For isolates tgbS1, tgbS2, tgbS4, and tgbS6, the starch hydrolysis test is positive. The presence of amylase enzymes is shown by the isolates’ ability to hydrolyze starch (Whitman and Parte 2009). Arabinose metabolism confirms the identity of IUBFASM_tgbS5 as it showed strong positive results with a yellow color, and IUBFASM_tgbS4 and IUBFASM_tgbS6 showed moderately positive results.

##### Citrate utilization

The isolates IUBFASM_tgbS1 and IUBFASM_tgbS2 showed strong positive color change, followed by moderate change by IUBFASM_tgbS4 and tgbS6, indicating utilization of citrate. Citric acid can be used as the exclusive carbon source by these isolates (Garcia 2010). IUBFASM_tgbS3 and tgbS5 gave negative results.

##### H_2_S and other gas production assay

None of the isolates generate H₂S from substances that include sulfur (Cappuccino and Sherman 2005). Only IUBFASM_tgbS5 showed the ability of gas production in the assay.

##### Oxidase and Catalase test

IUBFASM_tgbS1, tgbS2, tgbS4, and tgbS6 showed positive (+) in the Oxidase test, indicating Cytochrome c oxidase, a component of the electron transport chain, is present in these isolates (Versalovic 2011). Oxidase-negative bacteria rely on other pathways for energy production, such as fermentation or anaerobic respiration. All of the isolates had a positive (+) catalase test, meaning that catalase, which converts hydrogen peroxide into oxygen and water, is produced by all isolates (Cappuccino and Sherman 2005).

##### MR-VP test to differentiate glucose fermentation pathways

Methyl Red (MR) Test is moderately positive (+) for S3, showing a brownish appearance, and was found positive for S5. These isolates ferment glucose to yield stable acid end products. A positive Methyl Red (MR) test indicates that the bacterium has undergone mixed acid fermentation of glucose, resulting in the production of strong acids such as lactic acid, acetic acid, formic acid, and succinic acid. These acids lower the pH of the medium, which turns the pH indicator methyl red from yellow to red. The isolates IUBFASM_tgbS1, tgbS4, and tgbS6 gave positive (+) Voges-Proskauer (VP) Test. A positive VP test means that the bacterium predominantly ferments glucose via the butylene glycol pathway, which produces acetoin as a major end product.

(Versalovic 2011).

IUBFASM_tgbS5 is noteworthy since it tests positive for gas generation, glucose and lactose fermentation, indole formation, and MR, indicating that it might be a member of the Enterobacteriaceae family. Citrate is a valuable characteristic for distinction that S1, S4, and S6 can employ. The isolates appear to be from various groups or species based on their biochemical profiles, with S5 displaying the most unique characteristics.

#### 3.3.3. Profiling through 16S rRNA Sequencing

For correctly identifying the bacterial stain at the molecular level, 16S rRNA analysis was carried out. Based on the BLAST analysis with the NCBI 16S rRNA database, it was found that the microbes are from the *Bacillus, Stutzerimonas, Acinetobacter*, and *Enterobacteriaceae* genus (Table 2). From the 16S rRNA-based amplicon sequencing (supplementary figure 2), it was revealed that in IUBFASM_tgbS3 there might be a mixture of bacteria from both *Bacillus* and *Acinetobacter,* showing properties from both.

**Table 2:** Molecular Identification results of the isolates using 16S rRNA sequencing.

| ID of the Isolates | Practical plate ID | Genus from taxonomy | BLAST alignment match | Percent Identity | Bit score |
| --- | --- | --- | --- | --- | --- |
| IUBFAS M_tgbS1 | S1i3 | Bacillus | *Bacillus spizizenii* strain NRRL | 99.356 | 845 |
|  |  |  | *Bacillus subtilis* strain DSM 10 | 99.356 | 845 |
|  |  |  | *Bacillus inaquosorum* strain BGSC 3A28 | 99.356 | 845 |
|  |  |  | *Bacillus tequilensis* strain 10b | 99.356 | 845 |
| IUBFAS M_tgbS2 | S1i4 | Stutzerimonas | *Stutzerimonas stutzeri* strain DSM 5190 | 100 | 859 |
| IUBFAS M_tgbS3 | S2 | Bacillus | *Bacillus cereus* strain FDAARGOS_1084 | 100 | 861 |
|  |  |  | *Bacillus mycoides* strain JAS06 | 100 | 861 |
|  |  | Acinetobacter | *Acinetobacter baumannii* ATCC17978 | 99.785 | 856 |
| IUBFAS M_tgbS4 | S3i5 | Bacillus | *Bacillus spizizenii* strain NRRL | 100 | 841 |
|  |  |  | *Bacillus subtilis* strain DSM 10 | 100 | 841 |
|  |  |  | *Bacillus inaquosorum* strain BGSC 3A28 | 100 | 841 |
|  |  |  | *Bacillus tequilensis* strain 10b | 100 | 841 |
| IUBFAS M_tgbS5 | S4 | Enterobacteriaceae | *Escherichia coli* strain RHB34-CO2 | 100 | 859 |
|  |  |  | *Escherichia/Shigella* sp. Strain krb19-163 | 99.785 | 854 |
| IUBFAS M_tgbS6 | S3i4 | Bacillus | *Bacillus spizizenii* strain NRRL | 100 | 861 |
|  |  |  | *Bacillus subtilis* strain DSM 10 | 100 | 861 |
|  |  |  | *Bacillus inaquosorum* strain BGSC 3A28 | 100 | 861 |
|  |  |  | *Bacillus tequilensis* strain 10b | 100 | 861 |

#### 3.3.4 Biofilm Formation Ability of the Isolates

The isolate’s biofilm activity was evaluated by using a microtiter plate assay. The IUBFASM_tgbS3 isolate had the highest absorbance, followed by tgbS1 and tgbS6, indicating robust biofilm formation. IUBFASM_tgbS5 showed slightly lower biofilm formation compared to the other isolates. Biofilm formation pattern was similar among the isolates, but a bit reduced, when the assay was performed by incubating the isolates for 7 days, allowing production of biosurfactants (Figure 4). It indicates that the biosurfactants might have antibiofilm properties that hampered the biofilm formation ability in some isolates.

**Figure 4:**
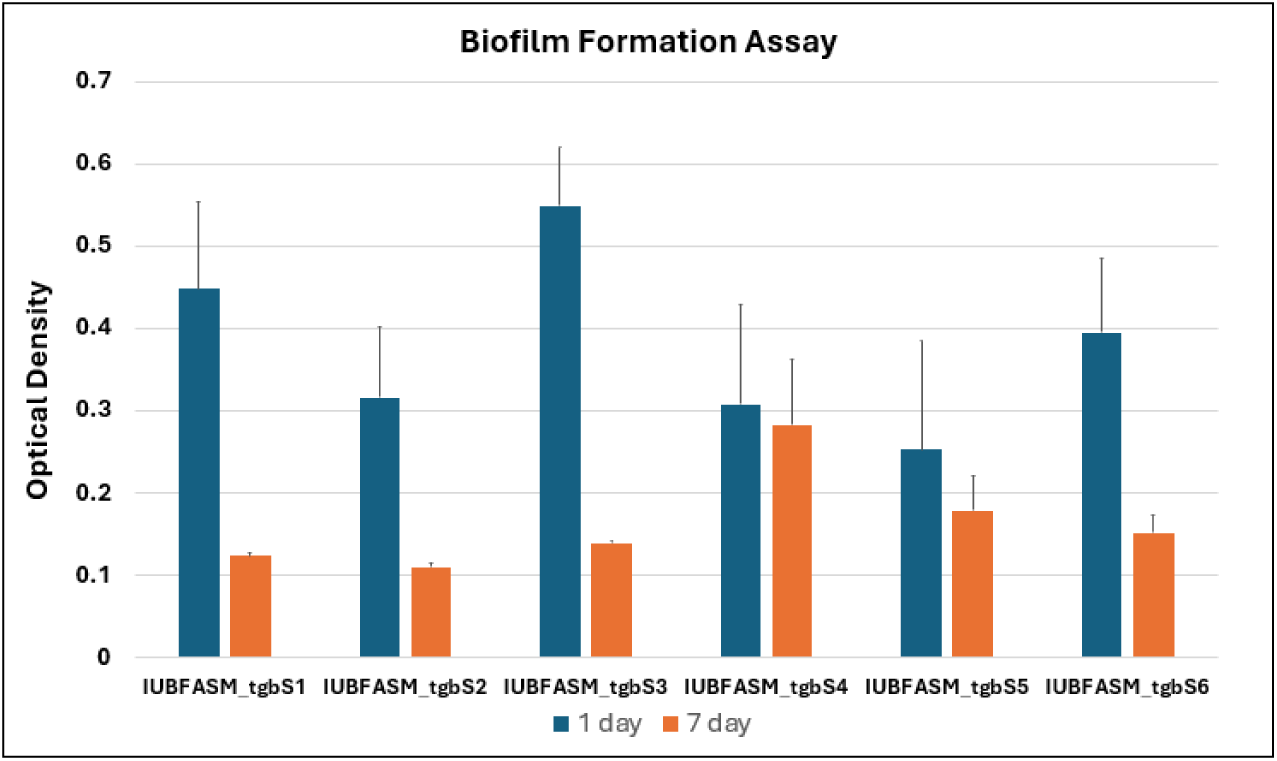
Biofilm Formation Ability of the isolates.

### 3.4 Characterization of the surfactant compound

#### 3.4.1 CTAB agar Test Reveals Ionic Nature of the Biosurfactant

In the case of CTAB method, a sign of ion matching between the cationic CTAB-methylene blue agar complex and anionic biosurfactant was the formation of a dark blue halo surrounding the culture (Saravanan and Vijayakumar 2012). IUBFASM_tgbS1 and IUBFASM_tgbS2 gave positive results, showing a dark blue halo. Since they interact with the cationic detergent CTAB, the surfactant is intended to be anionic and the blue zone was observed due to the presence of the methylene blue indicator. The other four isolates, however, showed moderate indication of blue color, but the lower intensity indicates the small yield of the biosurfactants in these bacteria.

**Figure 5:**
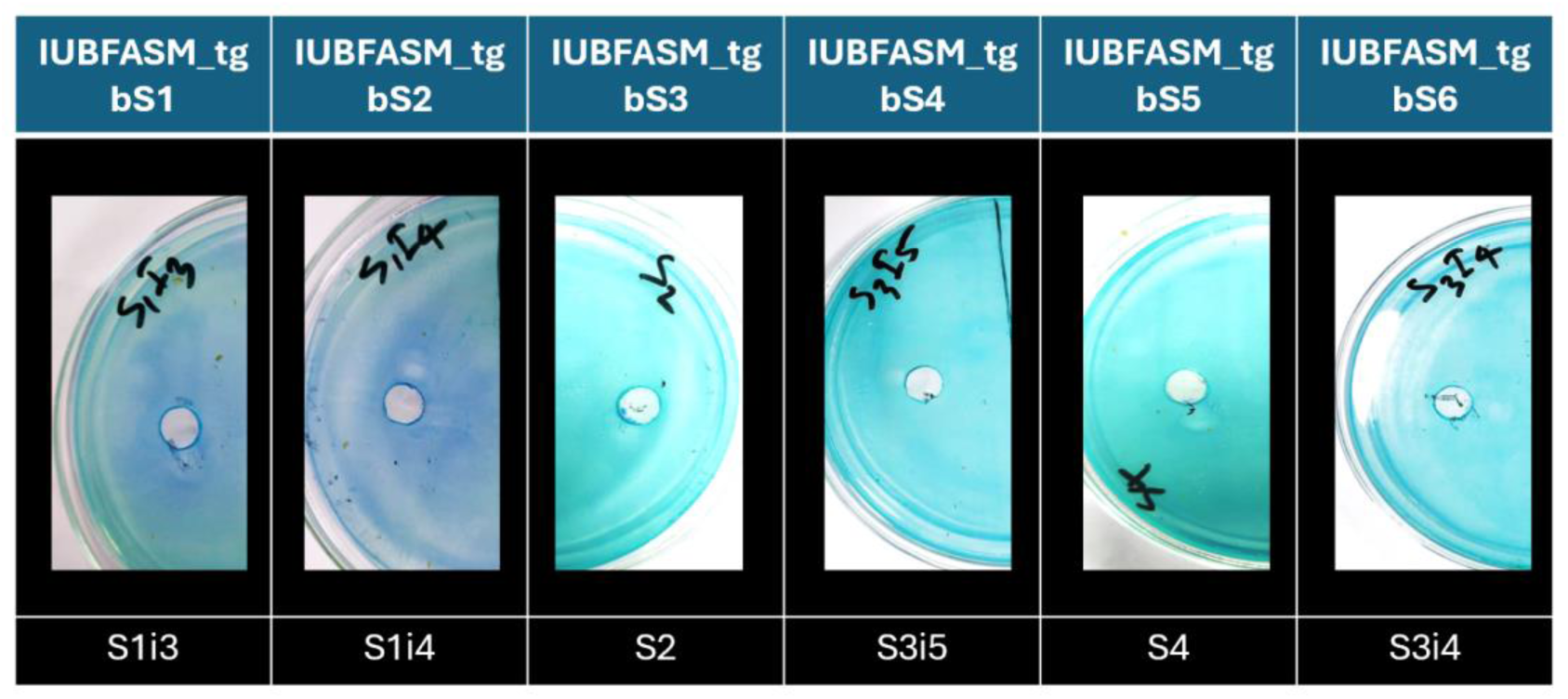
Biosurfactants showed a positive result at CTAB Methylene Blue agar complex

#### 3.4.2 Impact of pH, Temperature, and Salinity on Emulsification

The emulsion was found to be stable over a temperature range from 40°C to 100°C. For IUBFASM_tgbS2 and IUBFASM_tgbS6, the emulsion index increased from 15% and 8% at 40°C to 18.18% and 13% at 80°C, respectively. At 100°C, IUBFASM_tgbS2 showed highest stability of 44.44%. Except for IUBFASM_tgbS4, all isolates showed increased emulsification index at higher temperatures.

For all isolates, emulsion stability was found to be lowest under acidic conditions, with the exception of IUBFASM_tgbS1 and IUBFASM_tgbS6. IUBFASM_tgbS1, tgbS2, and tgbS6 showed stable emulsification index in both acidic and alkaline environments. The emulsion was found to be reasonably stable with the emulsion index in neutral conditions, although S1, S3, and S4 produced no results.

When the isolates were tested for emulsification index under 2%, 6% and 10% saline conditions, IUBFASM_tgbS6 showed tolerance to salt and exhibited a higher emulsification index at high NaCl salt conditions. IUBFASM_tgbS2 showed moderate E24% in 2% and 6% salt but not under 10%. IUBFASM_tgbS3 showed good E24% at 10% salt concentration, but not at the lower salt concentration.

Overall, higher temperature 80°C-100°C, and alkaline conditions (high pH) were found to be favourable for the isolates to generate higher emulsification. Among the six isolates, IUBFASM_tgbS6 showed promising stability of emulsification index in all pH, temperature, and saline conditions.

**Figure 6:**
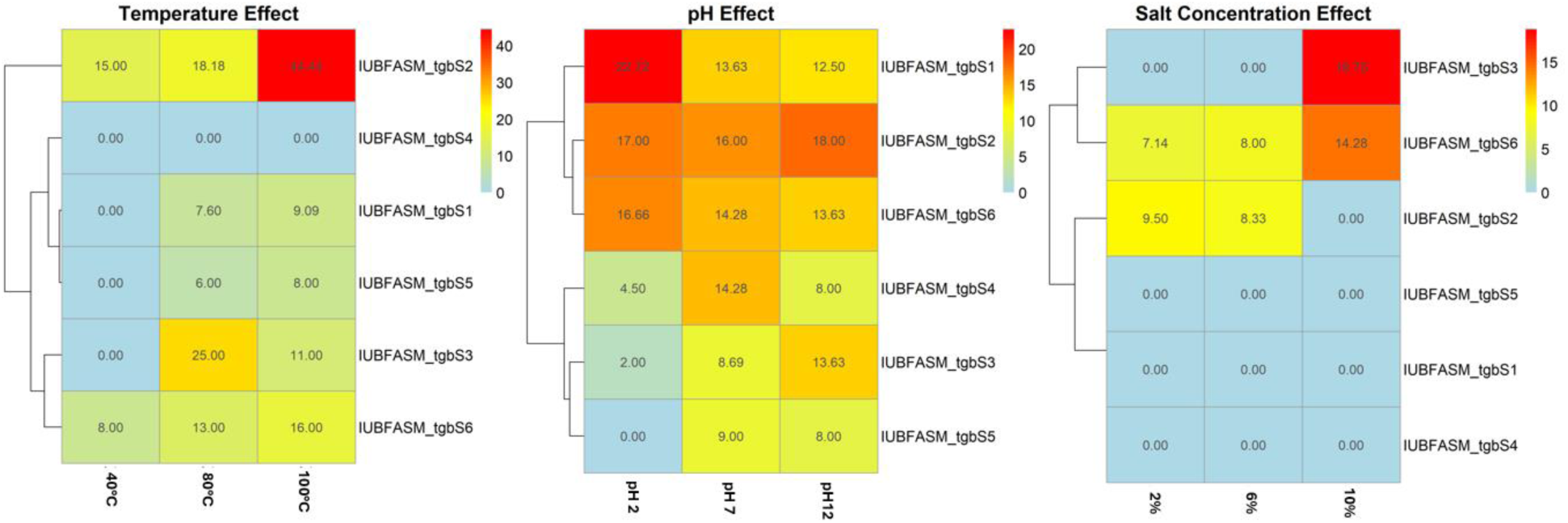
Emulsification Stability test

#### 3.4.3 Thin-layer chromatography and FTIR identified the lipopeptidic nature of the surfactants

After reacting with ninhydrin on the TLC plate, all of the isolates displayed a pink color, indicating that the biosurfactant contained peptide moieties and possibly lipopeptides. The sfactant compound of IUBFASM_tgbS2 was extracted and purified. A viscous, honey-colored product was obtained in small amount. Fourier Transform Infrared (FTIR) analysis revealed characteristic peaks corresponding to aliphatic chains (∼1450 cm-1), C-O stretching (∼1018 cm-1), and strong O-H stretching (∼3700-3900 cm-1), indicating a lipid-rich biosurfactant, possibly glycolipid in nature. *Stutzerimonas* can co-produce weak lipopeptides and glycolipids, which made a blurred FTIR signature and a weak amide band, and a strong lipid and sugar signal.

### 3.5 Additional Properties of the bacterial strains

#### 3.5.1 BATH assay measuring the adherence property of the surfactant

The BATH (Bacterial Adhesion to Hydrocarbons) test was used to determine the adherence % of the isolates, which evaluates cell hydrophobicity (Rosenberg et al. 1980). Since cells cling to oil droplets by generating surface-active biosurfactants, cell adhesion to hydrophobic substances like crude oil is regarded as an indirect technique to screen bacteria for biosurfactant synthesis. Among the isolates, IUBFASM_tgbS4 showed the highest adherence in the BATH assay, followed by IUBFASM_tgbS2. Lowest adherence was observed in IUBFASM_tgbS5, aligned with its other properties of biosurfactant production assays discussed above.

**Figure 7:**
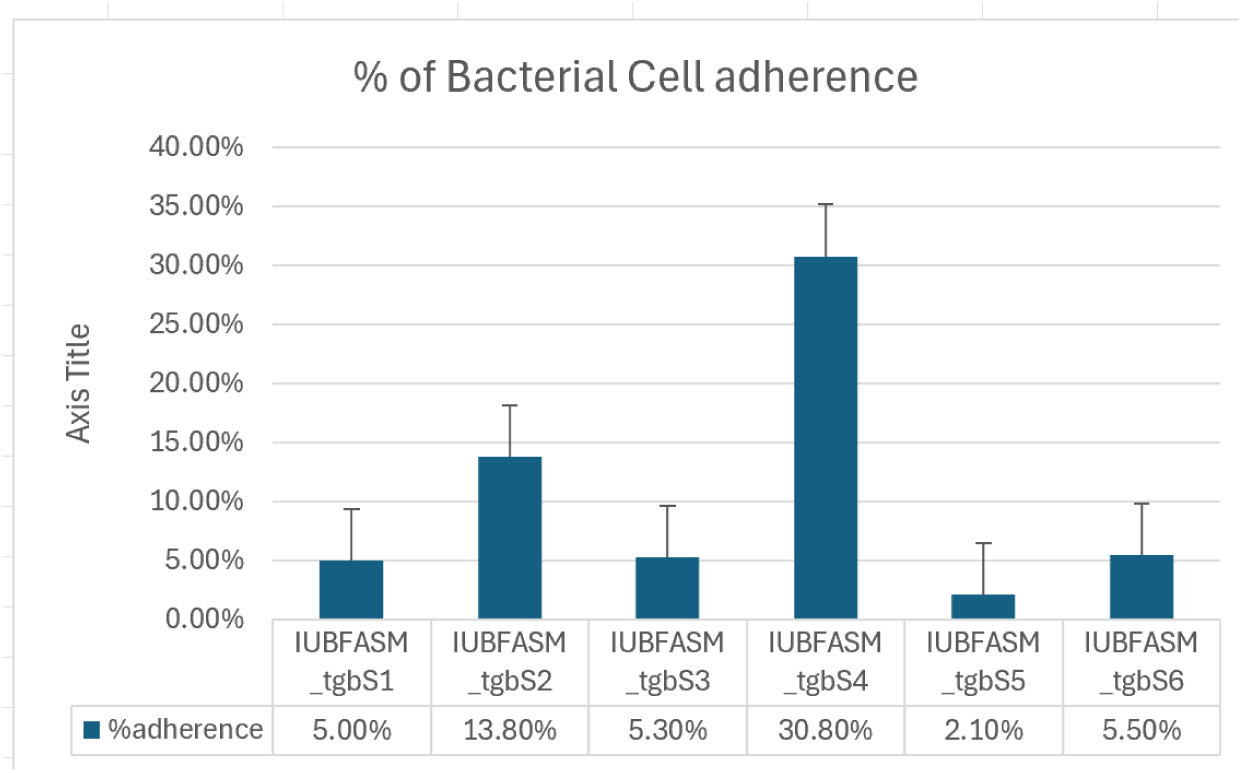
BATH assay

#### 3.5.2 Agar Diffusion Assay Demonstrated Antibacterial Effect

In case of antibacterial activity, all the isolates show possitive antibacterial activity against *Escherichia coli, Staphylococcus aureus, Vibrio cholerae Ogawa, Klebsiella,* and *Shigella* (Supplementary figure 3). Only IUBFASM_tgbS1 showed a negative result against *Staphylococcus aureus*.

#### 3.5.3 Isolates showed antifungal activity against plant pathogenic fungus

IUBFASM_tgbS1, tgbS2, tgbS4, and tgbS6 demonstrated antifungal activity in the well-diffusion assay against plant pathogenic fungus *Colletotrichum*, characterized by axenic inhibition zones free of satellite colonies or fungal encroachment. Both filtered and non-filtered cell-free extract was used to check whether the surfactant compound itself can combat fungus, or if it is the bacterial cells remaining in the non-filtered extract that are producing antifungal compounds to fight. IUBFASM_tgbS3 showed less activity for the non-filtered one, and IUBFASM_tgbS5 showed the opposite trend, where the non-filtered one gave good results. Except for filtered IUBFASM_tgbS5, tgbS6, and non-filtered IUBFASM_tgbS3, all isolates showed antifungal activity against *Colletotrichum*.

A reduced activity was observed against *Pestaletia*. Non-filtered IUBFASM_tgbS1 and IUBFASM_tgbS6 showed continued high inhibition. Filtered IUBFASM_tgbS2 and tgbS4 showed moderate activity. No activity was observed for IUBFASM_tgbS3 and tgbS5. Overall, all filtered extracts showed inhibition against *Colletotrichum* except IUBFASM_tgbS5 whereas IUBFASM_tgbS2 and tgbS4 filtered extracts showed inhibition against *Pestaletia*.

**Figure 8:**
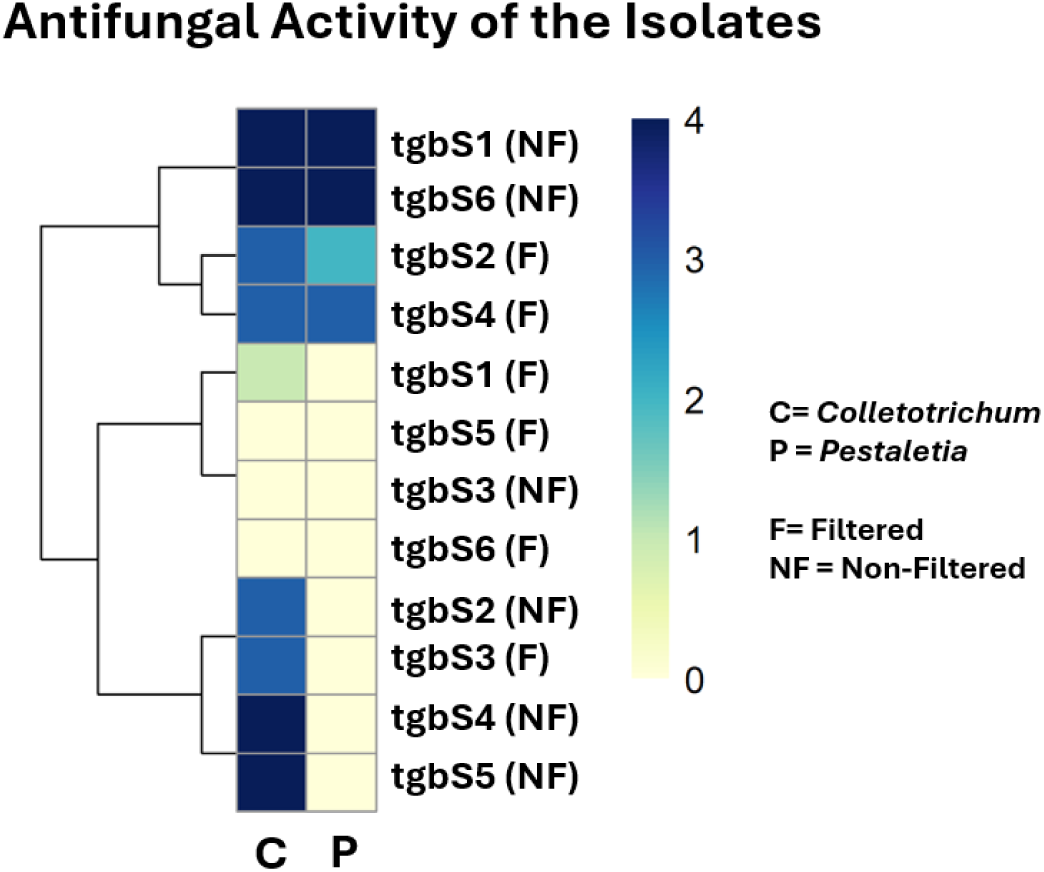
Antifungal Activity against plant pathogenic fungus

#### 3.5.4 Plant Growth Promotion Ability

Enhanced percentage of germination was observed in the *Oryza sativa* (indica) rice variety BRRI dhan28 upon application of the biosurfactant-producing bacteria. Even upon application of 150mM salt along with the bacteria, IUBFASM_tgbS1, tgbS2, and tgbS6 showed better germination. Highest germination was observed upon application of IUBFASM_tgbS4 and tgbS3, followed by tgbS5, without salt. IUBFASM_tgbS1 performed best to alleviate salt stress (Figure 9).

**Figure 9:**
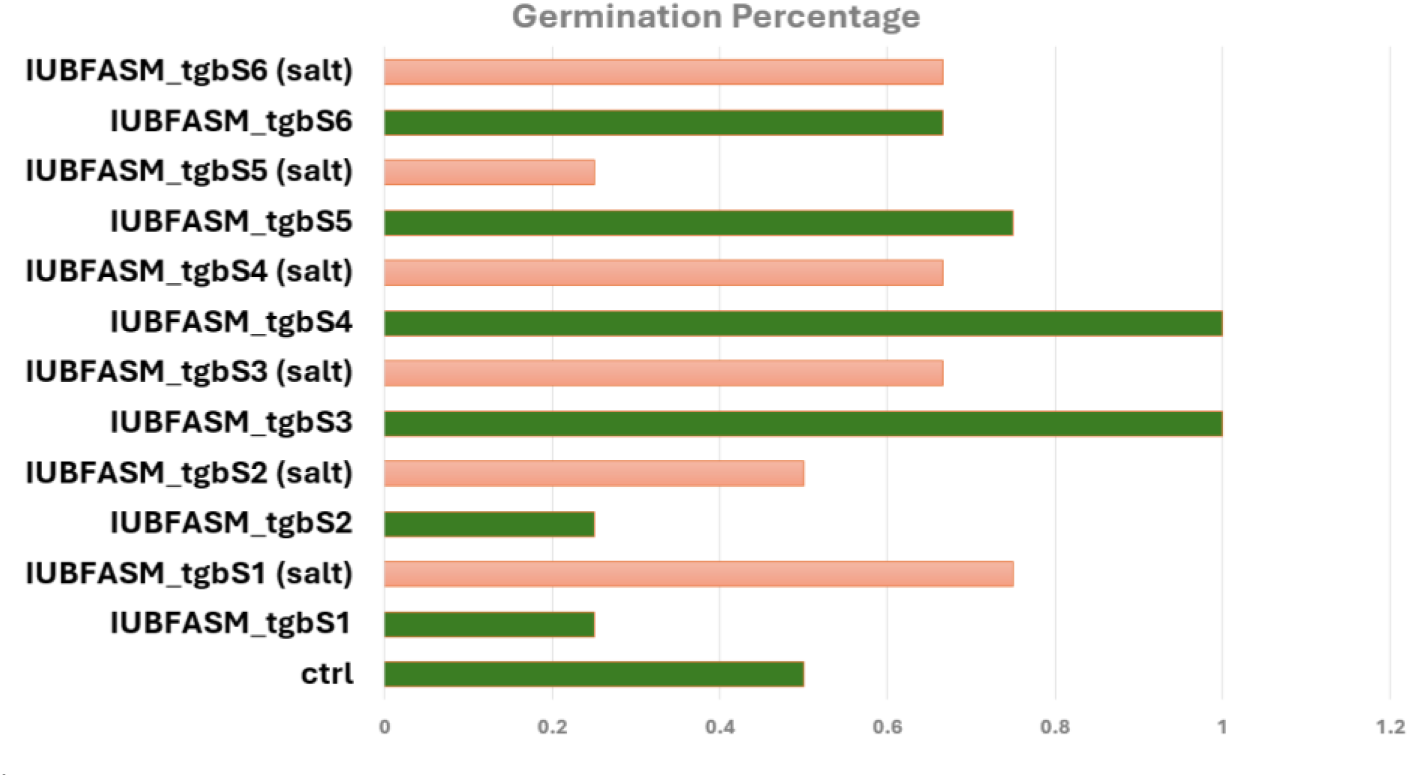
Germination Percentage of BRRidhan28 seeds upon application of a combination of biosurfactant-producing bacteria and salt

Growth promotion ability under salt stress was observed in all isolates except IUBFASM_tgbS6 (Figure 10). Tolerance to salinity was observed after 3 days. IUBFASM_tgbS3 and tgbs4 showed high growth promotion after 18 days, also (Supplementary figure 4). However, with only the cell-free extract, the plant growth promotion rate was not that good. Only IUBFASM_tgbS1 S1I3 under salt showed better performance (supp).

**Figure 10:**
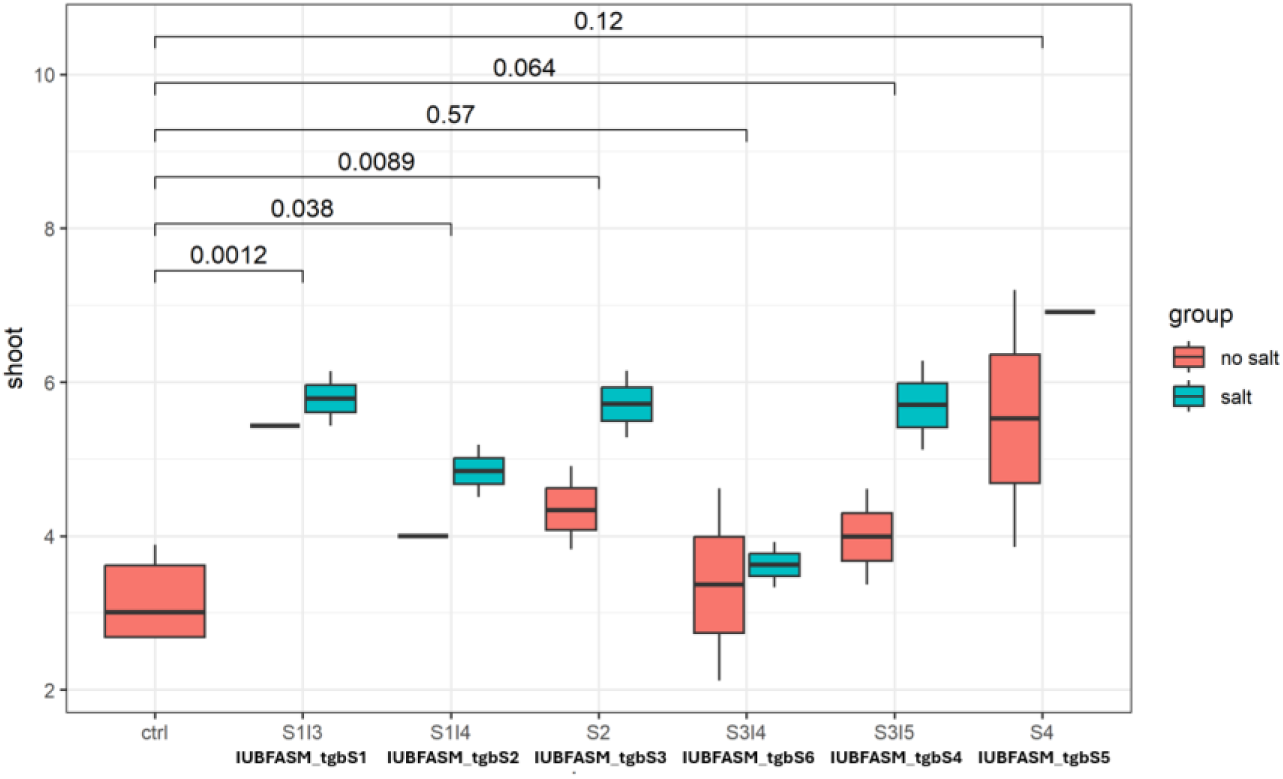
Plant growth promotion ability of the isolates after 3 days of salt stress

IAA test confirmed the production of Indole acetic acid by IUBFASM_tgbS2 and tgbS3, and strong IAA production by tgbS5 (Table 3). Gibberellic acid production was confirmed in all isolates, with IUBFASM_tgbS2 and tgbS4 exhibiting the highest levels, followed by tgbS6, tgbS3, and tgbS5. IUBFASM_tgbS1 exhibited the lowest GA production. Increased GA can help tolerate stress.

**Table 3:** Plant growth-promoting hormone production by the isolates.

|  | **IUBFASM**<br>**_tgbS1** | **IUBFASM**<br>**_tgbS2** | **IUBFASM**<br>**_tgbS3** | **IUBFASM**<br>**_tgbS4** | **IUBFASM**<br>**_tgbS5** | **IUBFASM**<br>**_tgbS6** |
| --- | --- | --- | --- | --- | --- | --- |
| **IAA** | - | + | + | - | ++ | - |
| **GA** | + | +++ | ++ | +++ | ++ | ++ |

Considering the germination percentage, seedling growth on petridish for 18 days, both with salt stress and without salt, aligns with the IAA and GA production ability of the isolates. It was confirmed that IUBFASM_tgbS3 and IUBFASM_tgbS4 showed high growth promotion ability.

## 4. DISCUSSION

The study reports the identification and isolation of six bacterial isolates with oil displacement property from industrial wastewater near Dhaka city, Bangladesh. These bacteria showed evidence of biosurfactant production in the oil displacement, drop collapse assay, and emulsification assay. Out of the six bacteria, three are from the *Bacillus* genus, one from *Stutzerimonus,* and another is from the *Enterobacteriaceae* group. One bacterial colony showed identity with both the Bacillus and Acinetobacter groups, reporting the possibility of mixed culture despite several attempts of isolation of pure culture. There are reports of coexistence of strains from these two genera in the environment, which can wok together towards biodegradation of pollutants (Yin et al. 2020). Identification of the bacterial consortium was confirmed through biochemical tests as well as sanger and ampliconseq based 16S rRNA sequencing, however ambiguity about some strains remained that cannot be overcome without multilocus typing that is beyond the scope of this report. In addition to hydrocarbon degradation ability, the isolates also showed antimicrobial activity as well as plant growth promotion ability, making them promising for not only industrial but also medical and agricultural applications. Proper identification of the bacteria and the characterization of the surfactant they produce are important for the utilization of these properties for industrial, medical, and agricultural usage.

### 4.1 Identification of the bacterial genera

For the identification of bacteria, the common molecular level method is sequencing of the 16S rRNA gene, which has 9 variable regions unique to different bacterial taxa. Using Sanger sequencing, all variable regions were covered to generate the sequence. Upon indication of mixed culture in one of the isolates 16S rRNA-based next-generation amplicon sequencing was performed, targeting the variable V3 and V4 region for confirmation of the results. S1i3 (IUBFASM_tgbS1), S3i5 (IUBFASM_tgbS4), and S3i4 (IUBFASM_tgbS6) showed the highest identity (99.35%) with 4 members of *Bacillus subtilis* species complex, i.e *Bacillus spizizenii, B. subtilis, B. inaquosorum* and *B. tequilensis.* In the biochemical assays, they were identified as positive in Gram staining, motility, oxidase, citrate, starch hydrolysis, and arabinose test. When we attempted to identify them more precisely, it appeared that, S1i3 (IUBFASM_tgbS1) can either belong to *Bacillus spizizeni*i or *Bacillus inaquosorum*. The isolates were able to grow in high pH conditions, characteristics relevant to *Bacillus spizizeni*i. Additionally, it exhibits comparatively lower PGPR activity while retaining a strong biofilm-forming capacity. On the other hand, the isolate showed less tolerance to acidic pH and showed low arabinose fermentation, exhibiting similarity with *Bacillus inaquosorum*. Without multilocus typing, revealing their true identity is not possible and beyond the scope of this report. S3i5 (IUBFASM_tgbS4) showed high antifungal activity, high adhesion in BATH assay, and less tolerance to extreme environments like high pH and temperature. It showed a good germination percentage under salt stress. These activities indicate it might be from the *Bacillus tequilensis* group. S3i4 (IUBFASM_tgbS6) showed better tolerance, high emulsification, and antimicrobial activity. Genome analysis of this strain categorized it in the *Bacillus subtills* group (Khatun and Elias 2025). *Bacillus subtilis* group normally produces surfactin-type biosurfactants, a lipopeptide. In addition to the oil-degrading ability, they show pharmacological effects like antibacterial, antifungal, ant-mycoplasma, antiviral, anti-thrombolytic, and anti-inflammatory activities (Zhen et al. 2023). All three bacillus isolates discussed above repeatedly showed negative glucose fermentation ability in the carbohydrate fermentation tests, although bacillus species normally show positive results in glucose metabolism. However, considering all other properties, this does not exclude them from being *Bacillus subtilis* species complex, as they may vary among strains, and weak or delayed reactions have been reported.

IUBFASM_tgbS2 (S1i4) showed 99.36% identity with *Stutzerimonas stutzeri* from the 16S rRNA result previously classified as *Pseudomonus stutzeri*. It is a promising bacterium for hydrocarbon bioremediation due to its high biodegradation efficiency and biosurfactant production. This species shows optimal crude oil degradation at pH 7–8 and 30–40 °C, achieving up to 84% degradation through the action of enzymes such as laccase, alcohol dehydrogenase, and alkane hydroxylase (Parthipan et al. 2017). *S. stutzeri* produces rhamnolipid-type biosurfactants that significantly reduce surface tension, enhance oil displacement, and achieve a high emulsification index (E24 = 51.3%), thereby increasing hydrocarbon bioavailability. Additionally, *Stutzerimonas spp.* exhibit strong surface attachment, biofilm formation, and heavy-metal tolerance mediated by biosurfactant chelation and extracellular polymeric substances, while posing a lower pathogenic risk than *Pseudomonas aeruginosa,* making them suitable for environmental and industrial applications (Varjani and Upasani 2017).

IUBFASM_tgbS3 (S2) showed 100% similarity with the *Bacillus cereus* strain FDAARGOS_1084 chromosome and 99.79% similarity with *Acinetobacter baumannii*. Taxonomic identification from the Qiime2 16s rRNA amplicon-seq analysis also indicated the abundance of both bacteria in the culture (supplementary figure 2). This raises the importance and the difficulties in isolating pure cultures from environmental samples. However, it also created scopes of studying the beneficial properties of both bacteria and taking precautions about any harmful effects when they are in a mixture. It showed moderate to low emulsification ability but high biofilm formation ability and plant growth promotion during germination. According to Wang et al. (2024), they produce surfactin. In our study, IUBFASM_tgbS3 showed high tolerance to salinity, alkaline pH, and high temperature. On the other hand, Acinetobacter is an aerobic, non-motile, gram-negative bacterium found in soil, freshwater, marine environments, as well as wastewater systems. *Acinetobacter bumannii* has shown the capacity to produce extracellular biosurfactants, predominantly composed of glycolipids, lipopolysaccharides, and complex polymeric compounds (Desai and Banat 1997; Rosenberg and Ron 1999; Mukherjee et al. 2006). These also show promising potential in agriculture as bio-stimulants and biocontrol agents by enhancing nutrient availability, promoting plant growth, and suppressing phytopathogens. Acinetobacter biosurfactants are also reported to be utilized in food processing, pharmaceutical, and enhanced oil recovery (Mukherjee et al. 2006). Usually, both *Bacillus cereus* and *Acinetobacter baumannii* show positive results in the citrate utilization test; however, in this experiment, it appeared slightly negative, which indicates some strains might utilize citrate weakly and can take more time, or there can be competition with other sources like glucose.

IUBFASM_tgbS5 (S4) shows positive fermentation of glucose and lactose, positive in MR, gas, catalase, and indole production; on the other hand, it exhibited a negative result in citrate and oxidase activity tests. It showed 100% similarity with *Escherichia coli* from the *Enterobacteriaceae* group. It also showed 99% identity with *Shigella*, but this was ruled out with the help of the arabinose test, which showed a positive result and a characteristic of *E. coli,* not *Shigella.* This bacterium showed high adherence, produced auxin and gibberellic acid, and demonstrated PGPR activity during plant growth screening. These properties establish it as a good root colonizer and potential biofertilizer candidate. It showed evidence of a limited amount of biosurfactant production. This might help the bacteria to colonize on the root, forming biofilm by reducing the surface tension. It can also help nutrient solubilization and have antimicrobial activity. Chigede et al. (2024) reported isolation of a bacterium with 98.29% 16srRNA identity with *E coli* strain, indicating that *E coli* can create a limited amount of biosurfactant, as observed in the current study. Biosurfactant production can improve adherence, biofilm formation, and environmental applications.

### 4.2 Independence of Oil Displacement and Emulsification was observed in Bacterial Isolates

Reduction of surface tension is the property of a surfactant. We observed differential indices of emulsification ability in diesel and kerosene. Emulsification (E24%) reflects the ability to form stable oil–water emulsions, which not only depends on the biosurfactant amount but also involves high molecular weight bio emulsifiers and EPS components like polysaccharides, proteins, and lipids. Hence, in the case of the *Stutzerimonus* strain, we observed a high emulsification index compared to others, but lower oil displacement ability. It showed an unusual amount of low biofilm formation but higher bacterial adherence ability. It can be a reason for observing a low level of biofilm in this strain, as the interaction of the biosurfactant with anti-biofilm property can hamper biofilm formation and depends on the concentration of the biosurfactant, as well as the nutrient availability and hydrodynamics, which can alter the behaviour.

The Bacterial Adherence to Hydrocarbons (BATH) assay is used to determine cell surface hydrophobicity, which is often influenced by biosurfactant production. The principle is based on the ability of bacterial cells to adhere to hydrocarbons (e.g., hexadecane, toluene, xylene), which correlates with their surface hydrophobicity. In early log phases, when biosurfactant production is low, high hydrocarbon adherence is expected, and in the late or stationary phases, low adherence to hydrocarbon should be observed. This scenario explains the high adherence of IUBFASM_tgbS2 but the low amount of biofilm production.

### 4.3 Biosurfactant-Producing Isolates Exhibit Potential for Applications in Agriculture and Medicine

The antifungal assay against 2 plant pathogenic fungi demonstrated the difference between filtered and non-filtered cell-free extracts in combating the fungal mycelia growth. The filtered cell-free extract does not contain bacterial cells; hence, it shows the ability of the extract to combat the fungus. The biosurfactant might disrupt microbial membranes via surface tension reduction and pore formation. Considering all properties, the IUBFASM_tgbS1, tgbS2, and tgbS6 were found with strong oil displacement ability, and IUBFASM_tgbS3 and tgbS5 were found to be plant growth promoters. IUBFASM_tgbS4 showed good growth promotion; however, under salt stress, it did not perform well, but the strong biosurfactant producers (tgbS1, tgbS2, tgbS6) did. This indicates the importance of the biosurfactants for combating salt stress. Biosurfactants reduce surface tension and improve water filtration around plant roots, ensuring water availability. Moreover, anionic biosurfactants can bind Na^+^ and other toxic ions, mitigating ionic stress on plants. They can also help other plant growth-promoting bacteria to adhere to the root surface and colonize the root by forming biofilm.

In addition to the antimicrobial properties of the strains, their biosurfactant-producing ability can be useful against antibiotic-resistant bacteria. By coating surfaces, biosurfactants reduce microbial colonization and can accelerate wound healing. This property can also prevent device-associated infection and improve drug delivery systems. Due to limited resources, the surfactant product of all strains could not be purified, but it opens the opportunity for more translational research in the future, utilizing these resources.

## 5. Conclusion

We have successfully isolated six bacterial strains exhibiting diverse levels of biosurfactant production from an industrial waste effluent river near Dhaka city using the oil enrichment strategy. Most of the isolates that demonstrated promising oil displacement and emulsification ability were identified as members of the *Bacillus subtilis* complex group, while one isolate belonged to the *Stutzerimonas* group. Two other bacterial isolates belonging to the *Enterobacteriaceae* and *Bacillus* groups showed comparatively lower biosurfactant production but exhibited better plant growth promotion ability. All isolates displayed more or less antibacterial and antifungal activity against plant pathogenic fungi. Identification and characterization of these hydrocarbon waste-adapted strains highlights their potential for eco-friendly applications in pollution mitigation. The antimicrobial and plant growth promotion ability of these strains suggests their potent usage in the medical and agricultural sectors. This study underscores the value of industrial waste-rich environments as reservoirs of metabolically versatile microorganisms holding significant potential for the identification of novel bacterial strains and unique bioactive compounds that enable microbial adaptation to extreme environmental conditions.

## Acknowledgement

The authors acknowledge the funding from the Independent University Bangladesh Sponsored Research grant 2021-SELS-05 and the Ministry of Science and Technology (MOST) SRG-252355 granted to the corresponding author. We are also grateful to Dr. Jebunnahar Khondakar, Department of Life Sciences, IUB, for sharing plant pathogenic fungi and Dr. Syesd Shifat K Ahmed for initially proposing the study. Special thanks to Lab technician Md. Khayrul Islam for supporting sample collection.

## Authors’ Contribution

UFTA = Performed the wet lab works and planned the experiments, assisted in writing, MM = Repeated the wet lab works and assisted in writing, SME = Designed the experiments and wrote the manuscript

## Highlights

- Isolated and characterized six bacterial strains with biosurfactant-producing ability at different levels, confirmed by oil displacement and emulsification assay.
- Antibacterial and antifungal properties of the strains were explored with potential medical applications.
- Plant growth-promoting ability of the isolates, with and without salt stress, was confirmed.

## REFERENCES

Ali, S. R. and B. Rajak (2013). "Screening and characterization of biosurfactants producing microorganism formnatural environment (whey spilled soil)." Screening 3(13): 53–58.

Arima, K., A. Kakinuma, et al. (1968). "Surfactin, a crystalline peptidelipid surfactant produced by Bacillussubtilis: Isolation, characterization and its inhibition of fibrin clot formation." Biochemical and biophysical research communications 31(3): 488–494.

Banat, I. M., A. Franzetti, et al. (2010). "Microbial biosurfactants production, applications and future potential." Applied microbiology and biotechnology 87(2): 427–444.

Burkholder, P. R., A. W. Evans, et al. (1944). "Antibiotic Activity of Lichens*." Proceedings of the National Academy of Sciences 30(9): 250–255.

Campos, J. M., T. L. M. Stamford, et al. (2019). "Characterization and application of a biosurfactant isolated from Candida utilis in salad dressings." Biodegradation 30(4): 313–324.

Cappuccino, J. G. and N. Sherman (2005). Microbiology: a laboratory manual, Pearson/Benjamin Cummings San Francisco.

Chigede, N., Z. Chikwambi, et al. (2024). "Isolation and characterization of biosurfactant-producing microbes isolated from the gastrointestinal system of broiler birds fed a commercial diet." Animal Biotechnology 35(1): 2263771.

Cooper, D. G. and B. G. Goldenberg (1987). "Surface-active agents from two Bacillus species." Applied and environmental microbiology 53(2): 224–229.

Datta, M. and I. Chattopadhyay (2024). "Applications of microbial biosurfactants in human health and environmental sustainability: a narrative review." Discover Medicine 1(1): 160.

Davies, P. J. (2004). Plant hormones: biosynthesis, signal transduction, action!, Springer Science & Business Media.

Deleu, M. and M. Paquot (2004). "From renewable vegetables resources to microorganisms: new trends in surfactants." Comptes Rendus Chimie 7(6-7): 641–646.

Desai, J. D. and I. M. Banat (1997). "Microbial production of surfactants and their commercial potential." Microbiology and Molecular Biology Reviews 61(1): 47–64.

Dunlap, L. and D. Beckmann (1988). "Soluble hydrocarbons analysis from kerosene / diesel." proceedings of the conference on Petroleum hydrocarbons and Organic chemicals in ground water 1988: Prevention, Detection and Restoration, Dublin, Ohio, National Water Well Association.

Femi-Ola, T., O. Oluwole, et al. (2015). "Isolation and screening of biosurfactant-producing bacteria from soil contaminated with domestic waste water." Br J Environ Sci 3: 58–63.

Garcia, L. S. (2010). Clinical microbiology procedures handbook, American Society for Microbiology Press.

Gordon, S. A. and R. P. Weber (1951). "Colorimetric Estimation of indoleacetic Acid." Plant Physiology 26(1): 192–195.

Gudiña, E. J., V. Rocha, et al. (2010). "Antimicrobial and antiadhesive properties of a biosurfactant isolated from Lactobacillus paracasei ssp. paracasei A20." Letters in applied microbiology 50(4): 419–424.

Jain, D., D. Collins-Thompson, et al. (1991). "A drop-collapsing test for screening surfactant-producing microorganisms." Journal of Microbiological Methods 13(4): 271–279.

Khatun, T. and S. M. Elias (2025). "Genome assembly and gene identification of biosurfactant-producing bacteria for environmental bioremediation." Briefings in Bioinformatics 26(Supplement_1): i46–i46.

Klindworth, A., E. Pruesse, et al. (2012). "Evaluation of general 16S ribosomal RNA gene PCR primers for classical and next-generation sequencing-based diversity studies." Nucleic Acids Research 41(1): e1–e1.

Makkar, R. S. and S. S. Cameotra (1997). "Biosurfactant production by a thermophilic Bacillus subtilis strain." Journal of Industrial Microbiology and Biotechnology 18(1): 37–42.

Md, F. (2012). "Biosurfactant: production and application." J Pet Environ Biotechnol 3(4): 124.

Mohan, P. K., G. Nakhla, et al. (2006). "Biokinetics of biodegradation of surfactants under aerobic, anoxic and anaerobic conditions." Water Res 40(3): 533–540.

Moldes, A. B., R. Paradelo, et al. (2011). "Ex situ treatment of hydrocarbon-contaminated soil using biosurfactants from Lactobacillus pentosus." Journal of agricultural and food chemistry 59(17): 9443–9447.

Morikawa, M., Y. Hirata, et al. (2000). "A study on the structure–function relationship of lipopeptide biosurfactants." Biochimica et Biophysica Acta (BBA)-Molecular and Cell Biology of Lipids 1488(3): 211–218.

Mukherjee, S., P. Das, et al. (2006). "Towards commercial production of microbial surfactants." TRENDS in Biotechnology 24(11): 509–515.

Mulligan, C. N. (2005). "Environmental applications for biosurfactants." Environ Pollut 133(2): 183–198.

Muthusamy, K., S. Gopalakrishnan, et al. (2008). "Biosurfactants: Properties, commercial production and application." Current Science 94(6): 736–747.

Parthipan, P., P. Elumalai, et al. (2017). "Biosurfactant and enzyme mediated crude oil degradation by Pseudomonas stutzeri NA3 and Acinetobacter baumannii MN3." 3 Biotech 7(5): 278.

Patten, C. L. and B. R. Glick (2002). "Role of *Pseudomonas putida* Indoleacetic Acid in Development of the Host Plant Root System." Applied and environmental microbiology 68(8): 3795–3801.

Rani, M., J. T. Weadge, et al. (2020). "Isolation and characterization of biosurfactant-producing bacteria from oil well batteries with antimicrobial activities against food-borne and plant pathogens." Front Microbiol 11: 64.

Rodrigues, L. R., J. A. Teixeira, et al. (2006). "Physicochemical and functional characterization of a biosurfactant produced by Lactococcus lactis 53." Colloids and Surfaces B: Biointerfaces 49(1): 79–86.

Rosenberg, E. and E. Z. Ron (1999). "High-and low-molecular-mass microbial surfactants." Applied microbiology and biotechnology 52(2): 154–162.

Rosenberg, M., D. Gutnick, et al. (1980). "Adherence of bacteria to hydrocarbons: a simple method for measuring cell-surface hydrophobicity." FEMS Microbiol lett 9(1): 29–33.

Rostás, M. and K. Blassmann (2009). "Insects had it first: surfactants as a defence against predators." Proceedings of the Royal Society B: Biological Sciences 276(1657): 633–638.

Saiyam, D., A. Dubey, et al. (2024). "Lipopeptides from Bacillus: Unveiling biotechnological prospects—Sources, properties, and diverse applications." Brazilian Journal of Microbiology 55(1): 281–295.

Salihu, A., I. Abdulkadir, et al. (2009). "An investigation for potential development on biosurfactants." Biotechnology and Molecular Biology Reviews 3(5): 111–117.

Samadi, N., N. Abadian, et al. (2007). "Biosurfactant production by the strain isolated from contaminated soil." J. Biol. Sci 7(7): 1266–1269.

Saravanan, V. and S. Vijayakumar (2012). "Isolation and screening of biosurfactant producing microorganisms from oil contaminated soil." J. Acad. Indus. Res 1(5): 264–268.

Sharma, M. and S. Kaur (2015). "Inhibitory potential of Lactobacillus species isolated fromfermented dairy products against Escherichia coli and Staphylococcus aureus." Journal of Gastrointestinal Infections 3: 45–50.

Siegmund, I. and F. Wagner (1991). "New method for detecting rhamnolipids excreted byPseudomonas species during growth on mineral agar." Biotechnology Techniques 5(4): 265–268.

Simpson, D. R., N. R. Natraj, et al. (2011). "Biosurfactant-producing Bacillus are present in produced brines from Oklahoma oil reservoirs with a wide range of salinities." Appl Microbiol Biotechnol 91(4): 1083–1093.

Singh, A., J. D. Van Hamme, et al. (2007). "Surfactants in microbiology and biotechnology: Part 2. Application aspects." Biotechnology advances 25(1): 99–121.

van der Vegt, W., H. C. van der Mei, et al. (1991). "Assessment of bacterial biosurfactant production through axisymmetric drop shape analysis by profile." Applied microbiology and biotechnology 35(6): 766–770.

Varjani, S. J. and V. N. Upasani (2017). "A new look on factors affecting microbial degradation of petroleum hydrocarbon pollutants." International Biodeterioration & Biodegradation 120: 71–83.

Versalovic, J. (2011). Manual of clinical microbiology, American Society for Microbiology Press.

Wang, X., J. Gao, et al. (2024). "Analysis of surfactant production by Bacillus cereus GX7 and optimization of fermentation conditions." Colloids and Surfaces B: Biointerfaces 233: 113629.

Whitman, W. B. and A. C. Parte (2009). Systematic bacteriology, Springer-Verlag New York.

Xia, W., Z. Du, et al. (2014). "Biosurfactant produced by novel Pseudomonas sp. WJ6 with biodegradation of n-alkanes and polycyclic aromatic hydrocarbons." Journal of Hazardous Materials 276: 489–498.

Yin, C.-F., Y. Xu, et al. (2020). "Biodegradation of polyethylene mulching films by a co-culture of Acinetobacter sp. strain NyZ450 and Bacillus sp. strain NyZ451 isolated from Tenebrio molitor larvae." International Biodeterioration & Biodegradation 155: 105089.

Zhen, C., X. F. Ge, et al. (2023). "Chemical structure, properties and potential applications of surfactin, as well as advanced strategies for improving its microbial production." AIMS Microbiol 9(2): 195–217.

